# Correcting mutant CFTR with a stabilizing nanobody reveals a novel active conformation of the channel

**DOI:** 10.1101/2025.10.08.681081

**Authors:** Marie Overtus, James N. Charlick, Tihomir Rubil, Andrew S. Paige, Blaine J. Loughlin, Zachary Rich, Mayuree Rodrat, Zhengrong Yang, Anita Balázs, John C. Kappes, Marcus A. Mall, David N. Sheppard, John F. Hunt, Cedric Govaerts

## Abstract

Defects in protein trafficking underlie many genetic diseases, including cystic fibrosis (CF), where the predominant F508del mutation destabilizes the cystic fibrosis transmembrane conductance regulator (CFTR) channel, leading to its degradation. To provide a protein-specific chaperone, we used lipid nanoparticles to deliver mRNA encoding T2a, a nanobody that thermally stabilizes CFTR via high-affinity binding to nucleotide-binding domain 1 (NBD1). When combined with clinically-approved correctors, T2a considerably improved F508del-CFTR maturation, plasma membrane expression, and channel activity. Single-channel recordings revealed that nanobody binding sustained channel activity by promoting both full open and sub-conductance gating states and protecting F508del-CFTR against thermal deactivation. Because T2a binding to NBD1 prevents the ATP-dependent NBD1-NBD2 association that drives canonical channel opening, the observation of channel activity in the presence of the nanobody suggested adoption of a novel open-channel CFTR structure. This inference was confirmed by cryo-EM analyses of CFTR in the presence of T2a, which revealed a novel open-channel conformation in which NBD1 binds to an alternative site on CFTR incompatible with NBD dimerization. Our findings establish a new paradigm to correct protein trafficking by stabilizing misfolded domains with targeted nanobodies and demonstrate a broadly applicable framework to treat CF and related protein misfolding diseases.

## INTRODUCTION

Cystic Fibrosis (CF) is the prototypical protein trafficking disease ^1^, where mutations cause processing, stability and functional defects leading to pathology. In CF, mutations in the cystic fibrosis transmembrane conductance regulator (CFTR) disrupt anion transport, which results in extensive tissue damage in vital organs, such as the respiratory airways and gastrointestinal system ^2,3^. CFTR is composed of two transmembrane domains (TMDs) connected to two nucleotide-binding domains (NBDs) by structured intracellular loops (ICLs) and separated by a regulatory (R) domain ^4^. Mutations that affect CFTR maturation and trafficking (class II mutations) ^5^ are found throughout the protein, but the most prevalent is the deletion of phenylalanine 508 (p.Phe508del; legacy: F508del) ^2^, located in NBD1, which causes thermal instability of the domain ^6,7^ that impairs assembly and maturation of the channel ^8,9^. Because the impact of F508del can be substantially suppressed either by compensating (stabilizing) mutations in NBD1 ^6,10,11^ or lowering the temperature during protein biogenesis ^12^, it is expected that molecular chaperones that stabilize NBD1 might be valuable therapeutics for people with CF bearing class II mutations ^13,14^.

The last decade has seen a revolution in the treatment of CF with the development of CFTR modulators that repair either protein trafficking (correctors) or channel gating (potentiators) ^3^. The coalescence of two correctors (elexacaftor and tezacaftor) with a potentiator (ivacaftor) in the triple combination therapy elexacaftor-tezacaftor-ivacaftor (ETI) ^15^ has had transformational benefits for most people with CF. While clinical outcomes are very positive overall, some studies have highlighted limitations of the treatment ^16–18^. Specifically, in people with CF and at least one F508del mutation, ETI restored CFTR activity to 40-50% of that in non-CF subjects ^16^, resembling the levels observed in non-classical CF ^19,20^. In addition, the response to ETI can vary within a patient cohort, with some individuals showing poor improvement, while others do not tolerate the therapy at all ^21^.

Structural studies demonstrate that CFTR modulators bind the channel at distinct locations within the TMDs ^22–24^ and more than 30 Å from F508 ^25^. Results from previous biophysical ^6,7^ and cell biological ^11,13,14^ studies suggest that molecules that specifically bind and stabilize NBD1 should rescue CFTR folding. Such molecules might provide alternative or complementary therapeutics for people with CF carrying the F508del mutation and potentially other class II mutations. We therefore reasoned that CFTR-specific nanobodies ^26,27^, which act as molecular chaperones, have therapeutic potential. For example, the nanobody T2a binds with high-affinity to both wild-type (WT) and F508del-CFTR, exerting a stabilizing effect of about 10-12 °C upon binding to either isolated NBD1 or full-length CFTR ^26^. However, the crystal structure of the NBD1-T2a complex indicated that binding of the nanobody should prevent NBD1:NBD2 dimerization and thus likely inhibit ATP-driven channel opening ^28^.

Here, we developed an mRNA LNP-based strategy to deliver a stabilizing nanobody to cells expressing F508del-CFTR and evaluated its ability to improve protein expression and function. We demonstrated that, by specifically stabilizing NBD1, the T2a nanobody dramatically increases the effects of correctors on F508del-CFTR protein expression and maturation. Using functional studies, we showed that this effect translates into a substantial synergistic improvement in channel activity at the plasma membrane. Cryo-EM studies of CFTR-T2a complexes demonstrated that the nanobody stabilizes at least two distinct conformations of the protein. In one state, CFTR is locked in a V-shaped inactive state in which T2a binding prevents the ATP-dependent NBD1-NBD2 association that drives canonical channel opening. However, remarkably, in the other state, the nanobody stabilizes a conformation where detachment of NBD1 enables pore formation in the absence of NBD1-NBD2 dimerization, revealing a novel active state of CFTR.

## RESULTS

### Synergistic correction of F508del-CFTR by stabilizing nanobody and CFTR correctors

To test the hypothesis that combining NBD1 stabilizers and the clinically-approved correctors should lead to synergistic rescue of F508del-CFTR, we selected nanobody T2a as a proof-of-concept NBD1-specific chaperone. However, for nanobody T2a to function as a folding chaperone, it must be delivered to the cell interior and be readily accessible to the newly synthesized polypeptide during the folding process. We used mRNA encapsulation into LNPs formulated for lung delivery ^29,30^ (see Methods) to deliver nanobody T2a ^26^ to the cell interior. Incubation of HEK293 cells with these LNPs achieved a transfection efficiency of at least 80%, leading to a dose-dependent expression of the nanobody (Figure S1).

**Figure S1.**
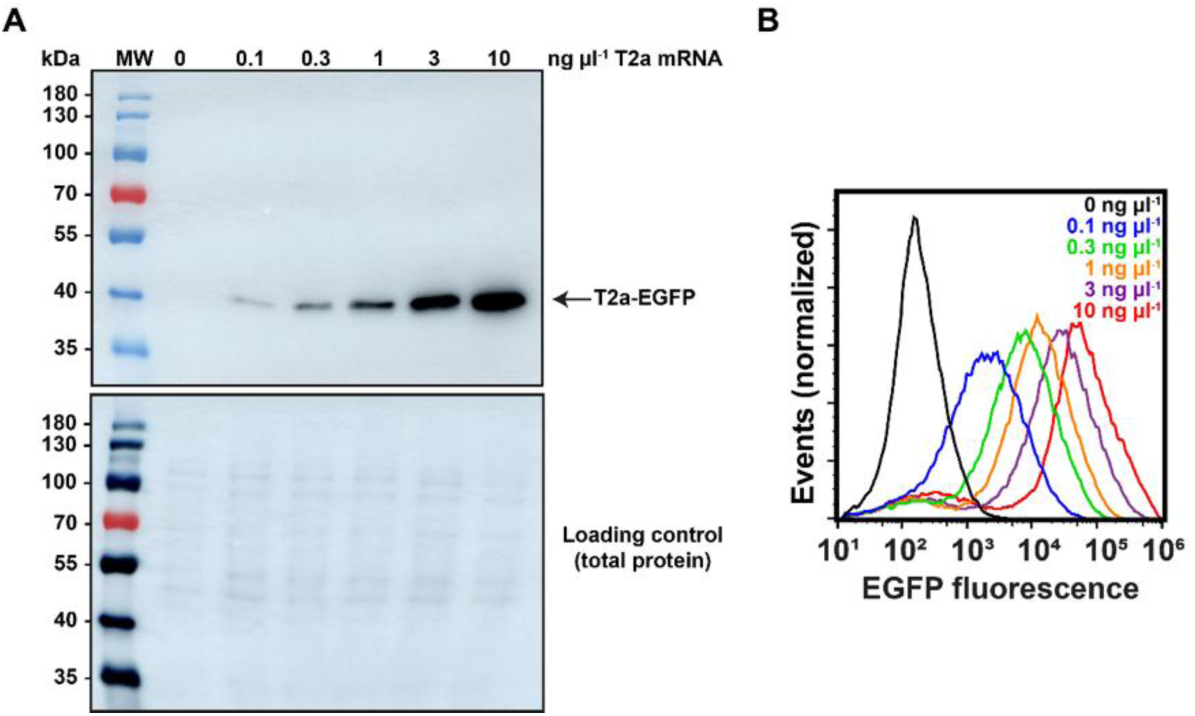
Transfection efficiency of T2a mRNA LNPs. (A, B) Dose-response relationship of T2a mRNA LNPs. (A) Top, immunoblot of cell lysates from HEK293 with T2a-EGFP detected with an anti-GFP antibody. Bottom, loading control with the total amount of protein detected using Cy5-NHS ester. (B) Flow cytometry analysis of T2a nanobody expression using the EGFP tag conjugated to the nanobody. The horizontal axis represents relative total cellular fluorescence, which is proportional to the functional expression of the T2a-EGFP fusion protein. Data are representative of at least 3 experiments.

### Combining approved correctors with nanobody expression substantially increases F508del-CFTR maturation

HEK293 and CF human bronchial epithelial cells (CFBE41o^−^) heterologously expressing human F508del-CFTR ^31^ were incubated with T2a LNPs in the absence or presence of ETI, and the efficiency of CFTR maturation was assessed by immunoblotting. Treatment with nanobody T2a alone led to little or no visible improvement in mature, fully glycosylated CFTR protein (band C), while ETI produced a variable but modest increase in mature CFTR protein in both cell lines (Figure 1A-D and Figure S2). By contrast, when the two treatments were combined, the increase in mature F508del-CFTR protein was considerably larger in both cell lines (Figure 1A-D and Figure S2). This result indicates that the combination treatment does not result in a simple additive effect but rather a strong synergy, supporting distinct mechanisms of action among the approved correctors and the stabilizing nanobody.

**Figure 1.**
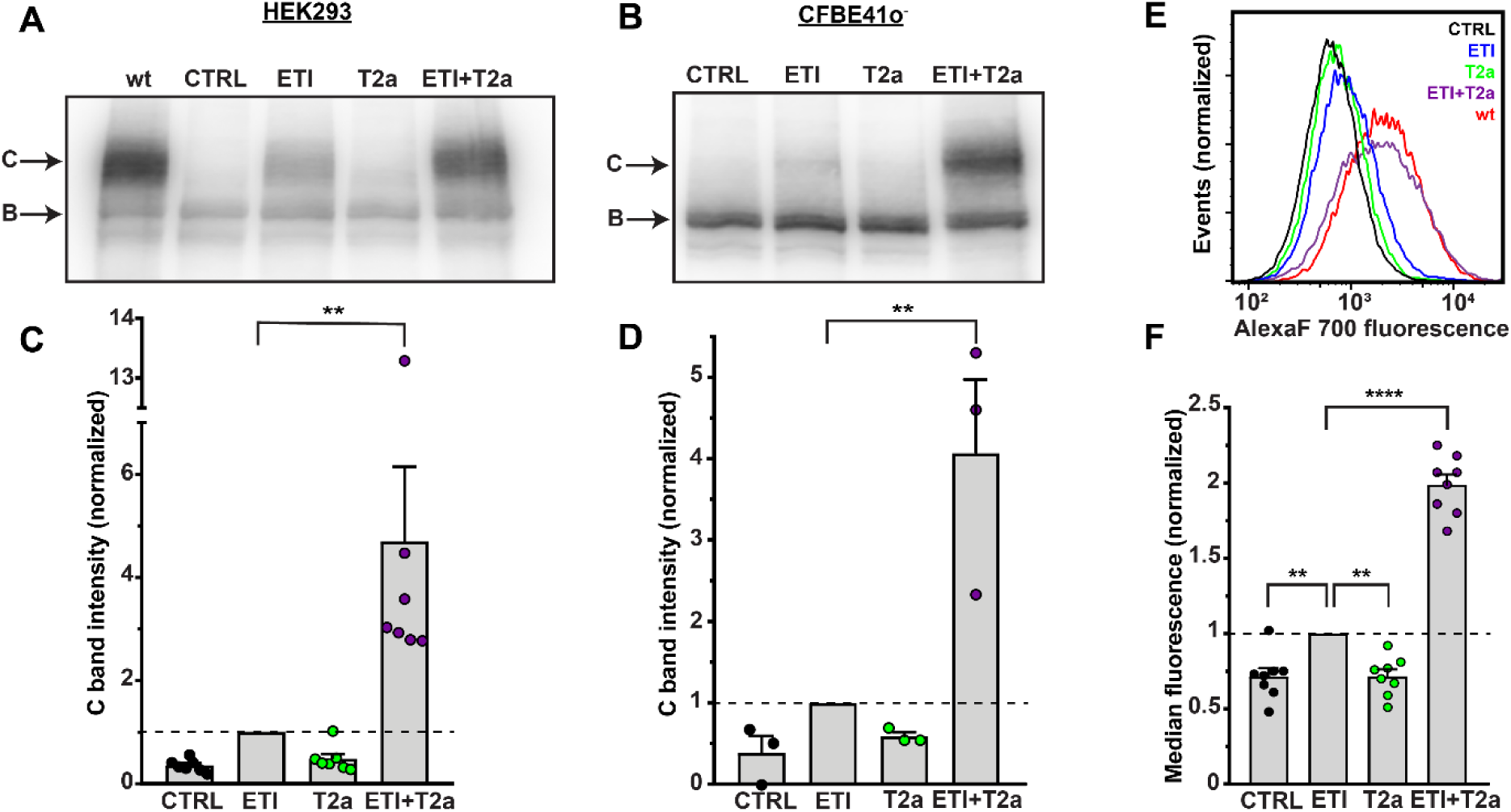
Nanobody T2a synergistically improves the efficacy of ETI-mediated F508del-CFTR maturation. (A, B) Immunoblots of WT- and F508del-CFTR maturation using cell lysates from CFTR-expressing HEK293 (A) and CFBE41o^−^ (B) cells treated with 0.3 ng µl^−1^ nanobody T2a mRNA LNP (T2a) either alone or together with 3 µM elexacaftor, 18 µM tezacaftor and 3 µM ivacaftor (ETI). Both the mature (band C) and immature (band B) CFTR protein were visualized with the anti-CFTR monoclonal antibody 596 that recognizes residues 1204-1211 of NBD2 ^32^. (C, D) Quantification of CFTR maturation by normalizing the data to ETI (C band intensity = 1; dashed lines). Symbols represent individual values and columns means ± SEM (n = 3-7); **, p < 0.01; one-way ANOVA. (E) Flow cytometry analyses of the effects of nanobody T2a mRNA LNPs and ETI treatment on F508del-CFTR expression at the plasma membrane of HEK293 cells. Cells were treated with T2a and ETI as described in A and B. An anti-HA antibody and a fluorescently labeled secondary antibody were used to detect the plasma membrane expression of F508del-CFTR with a genetically-encoded 3xHA tag in ECL4. For reference, the histogram of WT-CFTR (red) is also shown. Data were normalized to the number of events acquired in each condition. (F) Quantification of CFTR expression at the plasma membrane by normalizing the data as described in C and D. Symbols represent individual values and columns means ± SEM (n = 8); **, p < 0.01; ****p < 0.0001; one-way ANOVA.

**Figure S2.**
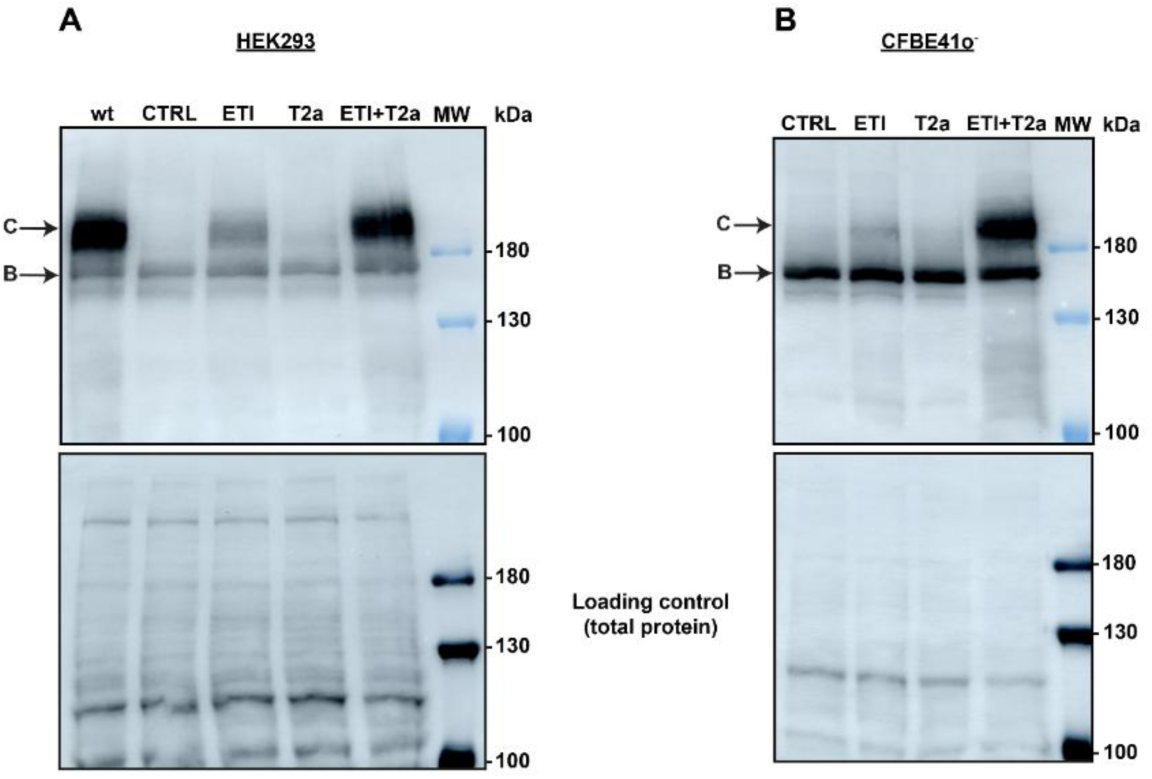
Immunoblots and loading controls for Figure 1A-B. Top, immunoblots of cell lysates from HEK293 (A) and CFBE41o^−^ (B) cells expressing CFTR. CFTR was detected using the anti-CFTR monoclonal antibody 596. Bands B and C represent the immature, core glycosylated and the mature, fully glycosylated CFTR protein, respectively. Bottom, loading controls of the immunoblots with the total amount of protein detected using Cy5-NHS ester.

As the HEK293 cells expressed F508del-CFTR with an engineered 3xHA tag in extracellular loop 4 (ECL4), we also examined the effects of the treatments on the plasma membrane expression of F508del-CFTR using flow cytometry by incubating cells with an anti-HA antibody and a fluorescent secondary antibody. Gating specifically on non-permeabilized cells, we observed that ETI modestly increased fluorescence, whereas nanobody T2a mRNA LNPs did not increase fluorescence above background (Figure 1E and F). By contrast, combining nanobody T2a with ETI yielded a dramatic increase in the plasma membrane expression of F508del-CFTR, reaching a fluorescence signal like that observed in HEK293 cells expressing WT-CFTR (Figure 1E). Furthermore, the distribution of fluorescence was monomodal, indicating that the combined treatment reached all cells.

### Increased maturation of F508del-CFTR upon nanobody treatment translates into enhanced functional rescue

To test whether improved F508del-CFTR maturation enhances the recovery of F508del-CFTR activity, we assessed CFTR function by halide-sensitive fluorescence quenching and the Ussing chamber technique. First, we assessed CFTR function with a fluorescence quenching assay using CFBE41o^−^ cells co-expressing F508del-CFTR and a halide-sensitive yellow fluorescent protein (YFP) ^33,34^.

After activating and potentiating F508del-CFTR with forskolin and ivacaftor, addition of iodide to the extracellular medium decreased YFP fluorescence in CFBE41o^−^ cells incubated with either nanobody T2a mRNA LNPs or elexacaftor and tezacaftor when compared to untreated cells with the correctors mediating the larger decrease (Figure 2A and B). Consistent with our CFTR maturation data (Figure 1), incubating CFBE41o^−^ cells co-expressing F508del-CFTR and YFP with elexacaftor, tezacaftor and nanobody T2a mRNA LNPs led to a much stronger decrease in YFP fluorescence (Figure 2A and B), demonstrating that the synergistic increase of mature F508del-CFTR protein at the plasma membrane confers greater channel activity.

**Figure 2.**
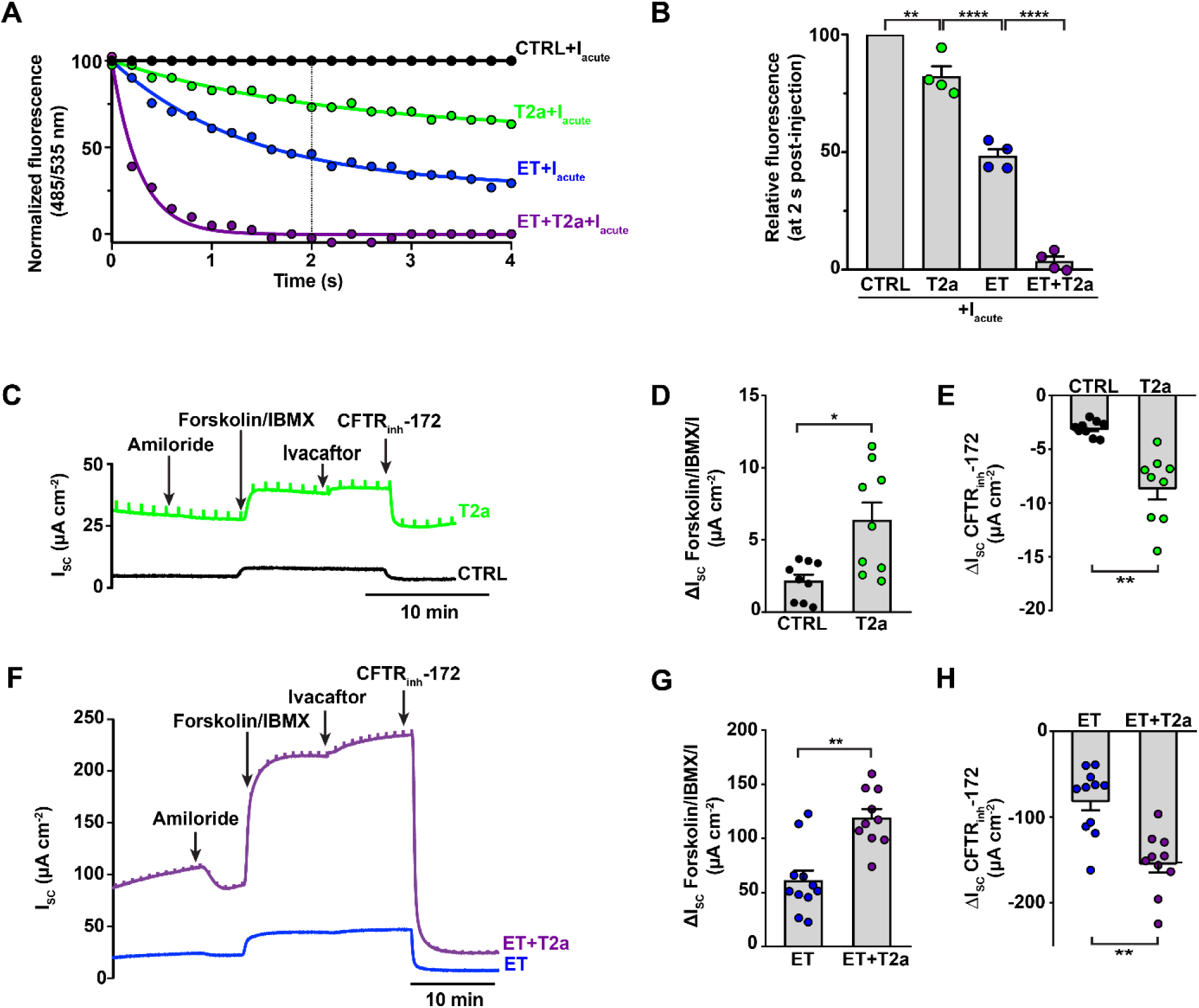
Nanobody T2a mRNA LNPs increase F508del-CFTR activity and synergize with ETI. (A) Halide sensitive-YFP quenching assay to measure F508del-CFTR activity in CFBE41o^−^ cells co-expressing F508del-CFTR and halide-sensitive YFP. The curves show YFP fluorescence quenching over the first 4 s following iodide injection after treating cells with 10 µM forskolin and 3 µM ivacaftor (I). Cells were incubated with (i) 2 ng µl^−1^ nanobody T2a mRNA LNPs (T2a), (ii) 3 µM elexacaftor and 18 µM tezacaftor (ET), (iii) ET and T2a (ET+T2a) or were untreated (CTRL). (B) Average YFP fluorescence 2 s after iodide injection for the conditions shown in A. Symbols represent individual values and columns are means ± SEM (n = 4); **, p < 0.01; ****, p < 0.0001; one-way ANOVA. (C) and (F) Representative I_sc_ recordings in CFBE41o^−^ cells expressing F508del-CFTR. Prior to the recordings, CFBE41o^−^ cells were incubated with either (i) 10 ng µl^−1^ nanobody T2a mRNA LNPs (T2a), (ii) 3 µM elexacaftor and 18 µM tezacaftor (ET), (iii) ET and T2a (ET+T2a) or were untreated (CTRL). Arrows indicate the sequential and cumulative addition of amiloride (100 µM), forskolin (10 µM)/IBMX (100 µM), ivacaftor (5 µM) and CFTR_inh_-172 (20 µM) to the apical solution with forskolin/IBMX also added to the basolateral solution. (D), (E), (G) and (H) Quantification of CFTR-mediated forskolin/IBMX/ivacaftor-induced I_sc_ (D and G) and CFTR_inh_-172-sensitive I_sc_ (E and H). Symbols represent individual values and columns are means ± SEM (n = 3-9); *, p < 0.05; **, p < 0.01; one-way ANOVA.

Finally, to evaluate the action of nanobody T2a on CFTR function at the tissue level, we measured cAMP-stimulated Cl^−^ currents in polarized F508del-CFTR-expressing CFBE41o^−^ epithelia with the Ussing chamber technique. CFTR-mediated Cl^−^ currents were quantified for untreated (CTRL) epithelia and those treated with (i) nanobody T2a, (ii) the correctors elexacaftor and tezacaftor, and (iii) correctors and T2a by measuring the change in short-circuit current (I_sc_) stimulated by cAMP-dependent activation with forskolin/3-isobutyl-1-methylxanthine (IBMX), potentiated by ivacaftor and inhibited with the CFTR inhibitor CFTR_inh_-172. Paralleling the results observed with the halide-sensitive fluorescence quenching assay (Figure 2A and B), treatment of F508del-CFTR-expressing CFBE41o^−^ cells with nanobody T2a mRNA LNPs modestly, but significantly, enhanced CFTR-mediated I_sc_ compared with untreated controls, while the correctors elexacaftor and tezacaftor again elicited greater CFTR-mediated I_sc_ than T2a-LNPs (Figure 2C–H). By contrast, combining T2a-LNPs with the correctors doubled the magnitude of CFTR-mediated I_sc_ compared to cells treated with the correctors alone (Figure 2F–H). Thus, the nanobody T2a robustly rescues F508del-CFTR expression and function in synergy with clinically-approved correctors, suggesting that it might have therapeutic potential.

### T2a modulates CFTR channel activity at the single-molecule level

To investigate whether T2a modulates the molecular behavior of CFTR, we used excised inside-out membrane patches from BHK cells heterologously expressing wild-type (WT) and F508del-CFTR. To magnify channel openings, we imposed a large Cl^−^ concentration gradient across membrane patches and clamped voltage at –50 mV. In the absence of nanobody, following phosphorylation with PKA, WT-CFTR displayed a bursting pattern of channel gating with openings interrupted by brief closures, separated by longer closures between bursts (Figures 3A, 3B and S3A), with a stable open probability (P_o_) of 0.3 ± 0.02 (n = 4) in the presence of ATP (0.3 mM). By contrast, the addition of 1 μM of purified T2a to the intracellular solution imposed a clear modal gating pattern on WT-CFTR (Figure 3A and 3B), which alternated between long (>2 s) inactive periods (P_o_ = 0 ± 0; n = 4) and active periods (P_o_ = 0.4 ± 0.03; n = 4) with little change in overall P_o_. We hypothesized that the inactive periods might represent the conformation where nanobody binding prevents NBD1:NBD2 dimerization ^26^ and, thus, channel opening ^28^ while the active periods might correspond to either nanobody unbinding or an active conformation that is compatible with both T2a binding and channel opening (see below). Importantly, the concentration of T2a used in these experiments (1 μM) was more than 2 orders of magnitude greater than the apparent affinity of this nanobody for either isolated NBD1 or full-length CFTR ^26^.

**Figure 3.**
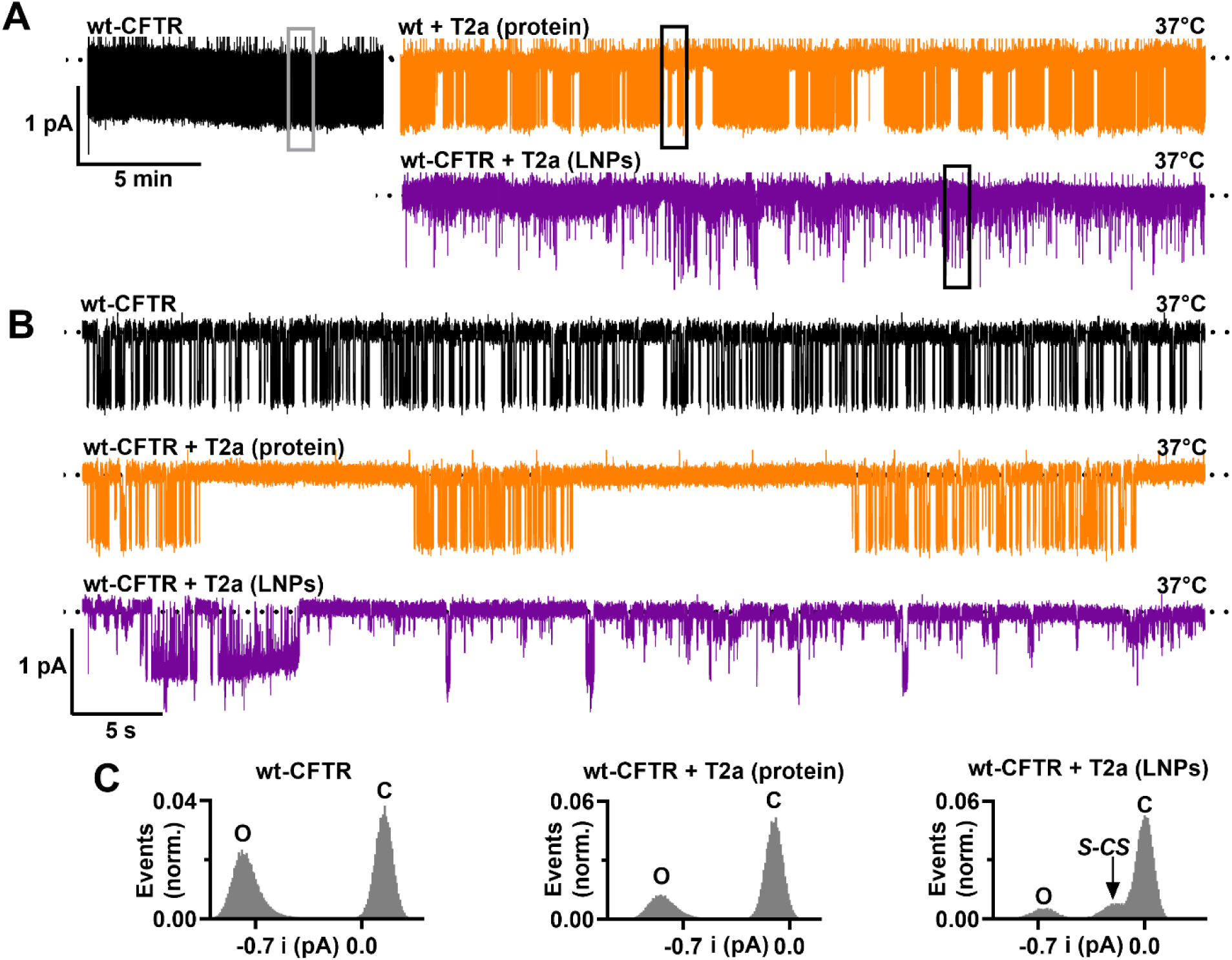
Nanobody T2a modifies the gating behavior of WT-CFTR. (A) Representative prolonged recordings of single WT-CFTR Cl^−^ channels in excised inside-out membrane patches from BHK cells stably expressing WT-CFTR acquired either in the absence (black) and presence of purified nanobody T2a (1 μM) in the intracellular solution (orange) or from a WT-CFTR-expressing BHK cell treated with T2a nanobody mRNA LNP (2 ng μl^−1^) for 24 h at 37 °C. ATP (wt and wt+T2a (protein), 0.3 mM; wt+T2a (LNPs), 1 mM) and PKA (75 nM) were continuously present in the intracellular solution; temperature was at 37 °C. Dotted lines indicate where channels are closed and downward deflections correspond to channel openings. (B) 60-s portions of the recordings shown in A taken from the boxed regions displayed on an expanded time scale. (C) Single-channel current amplitude histograms of the recordings shown in B; abbreviations: C, closed; O, open; S-CS, sub-conductance state.

**Figure S3.**
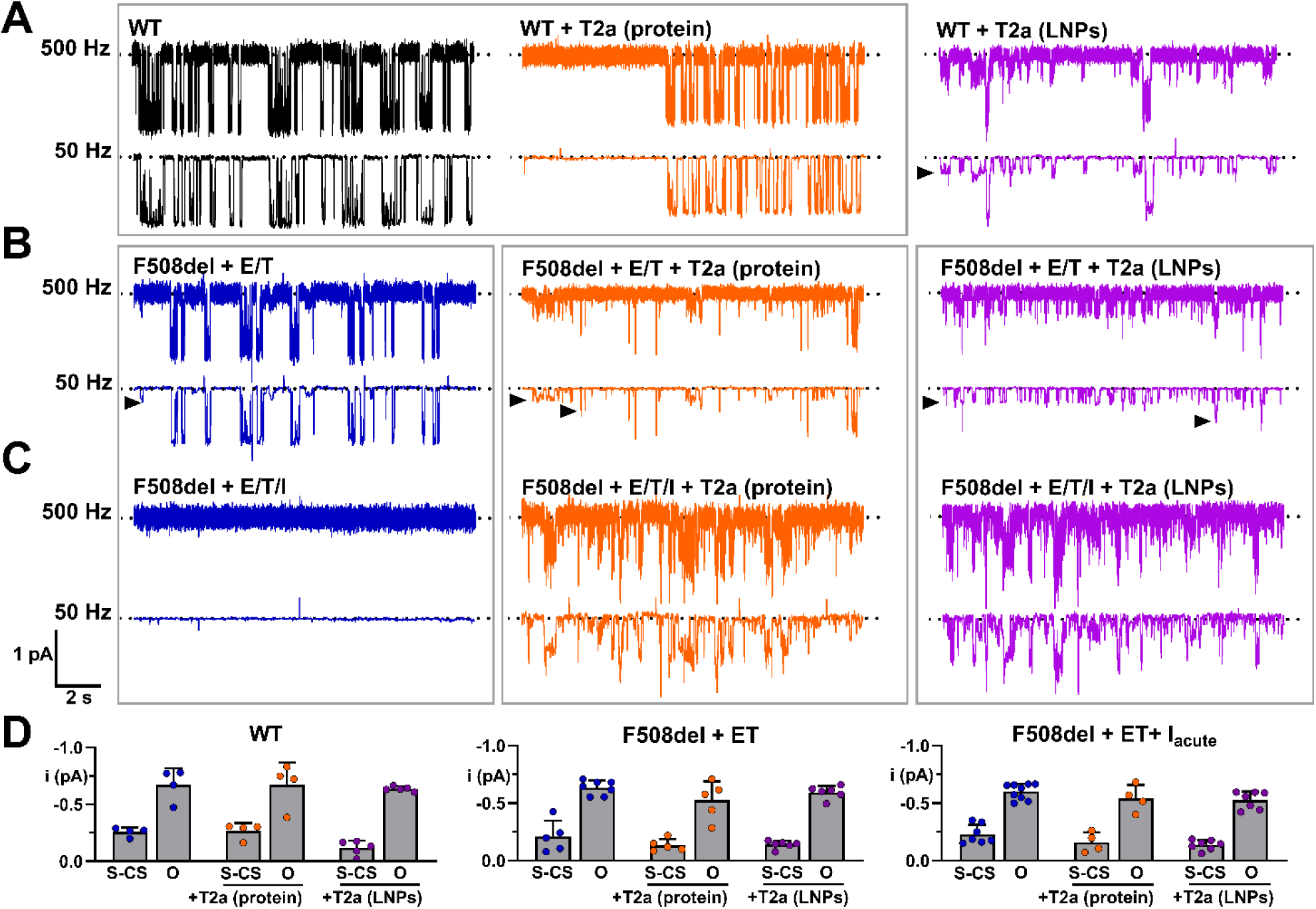
Nanobody T2a promotes opening of WT and F508del-CFTR Cl^−^ channels to sub-conductance states. Representative 12-s single-channel recordings of wild-type (A) and F508del-CFTR (B, C) in excised inside-out membrane patches that were either filtered at 500 Hz or additionally filtered at 50 Hz. The recordings were acquired either in the absence of nanobody, in the continuous presence of nanobody T2a (1 μM) in the intracellular solution or using membrane patches excised from cells treated with T2a nanobody mRNA LNPs (2 ng μl^−1^) for 24 h at 37 °C. Grey boxes identify recordings from the same membrane patches. To enhance the plasma membrane expression of F508del, F508del-CFTR-expressing BHK cells were treated with elexacaftor (E, 2 μM) and tezacaftor (T, 3 μM) for 24 h at 37 °C. Following channel activation by PKA (75 nM) and ATP (wt and wt+T2a (protein), 0.3 mM; wt+T2a (LNPs), 1 mM; F508del, 1 mM), which were then continuously present in the intracellular solution, ivacaftor (I, 1 μM) was added to the intracellular solution. The recordings in C were acquired >10 min after ivacaftor was added to the intracellular solution (see Fig. 4A – C). In A and B, arrowheads denote channel openings to sub-conductance states. (D) Quantification of the single-channel current amplitudes of the sub-conductance (S-CS) and full open (O) states for the different experimental conditions. Symbols represent individual values and columns are means ± SEM (wt, n = 4-5; F508del, n = 4-9).

However, when membrane patches were excised from WT-CFTR-expressing BHK cells incubated with T2a-LNPs, we observed a gating pattern distinct from that seen with acute addition of the purified nanobody. Figure 3A and 3B reveal that channel gating was highly variable, with transitions to the full open-state separated by numerous openings to small-amplitude sub-conductance states (S-CSs) and no prolonged closures like those observed when WT-CFTR was acutely treated with purified T2a. To resolve openings of CFTR to small-amplitude S-CSs, we digitally filtered single-channel recordings at 50 Hz (Figure S3), an approach that we have previously used to resolve openings of mouse CFTR to a tiny S-CS ^35^. Quantification of openings to S-CSs after low-pass filtering highlights the dramatic changes in single-channel behavior of WT-CFTR observed upon incubation with T2a LNPs. Figure 3C and S3D demonstrate a large reduction in openings to the full open-state and a marked increase in transitions to S-CSs. Thus, long-term incubation with the nanobody leads to different single-channel gating behavior.

We then investigated the effects of nanobody T2a on the single-channel behavior of F508del-CFTR rescued with the correctors elexacaftor and tezacaftor. When cells were treated with elexacaftor and tezacaftor only, F508del-CFTR showed moderate channel activity that was increased about two-fold by ivacaftor (Figure 4A). However, potentiation of F508del-CFTR by ivacaftor was short-lived, with current decreasing to background levels after 10-15 min (Figure 4A) in line with early observation that the potentiator has a destabilizing effect on mutant CFTR ^36^. By contrast, when similar measurements were performed in the presence of nanobody T2a, either as purified protein added to the intracellular solution (Figure 4B) or after incubation of F508del-CFTR-expressing cells with T2a LNPs (Figure 4C), the activity of F508del-CFTR was sustained for the entire duration of prolonged recordings (≥ 30 min) at 37 °C even when channels were potentiated by ivacaftor. Figure 4C demonstrates that this sustained activity of F508del-CFTR was inhibited by the CFTR inhibitor CFTR_inh_-172.

**Figure 4.**
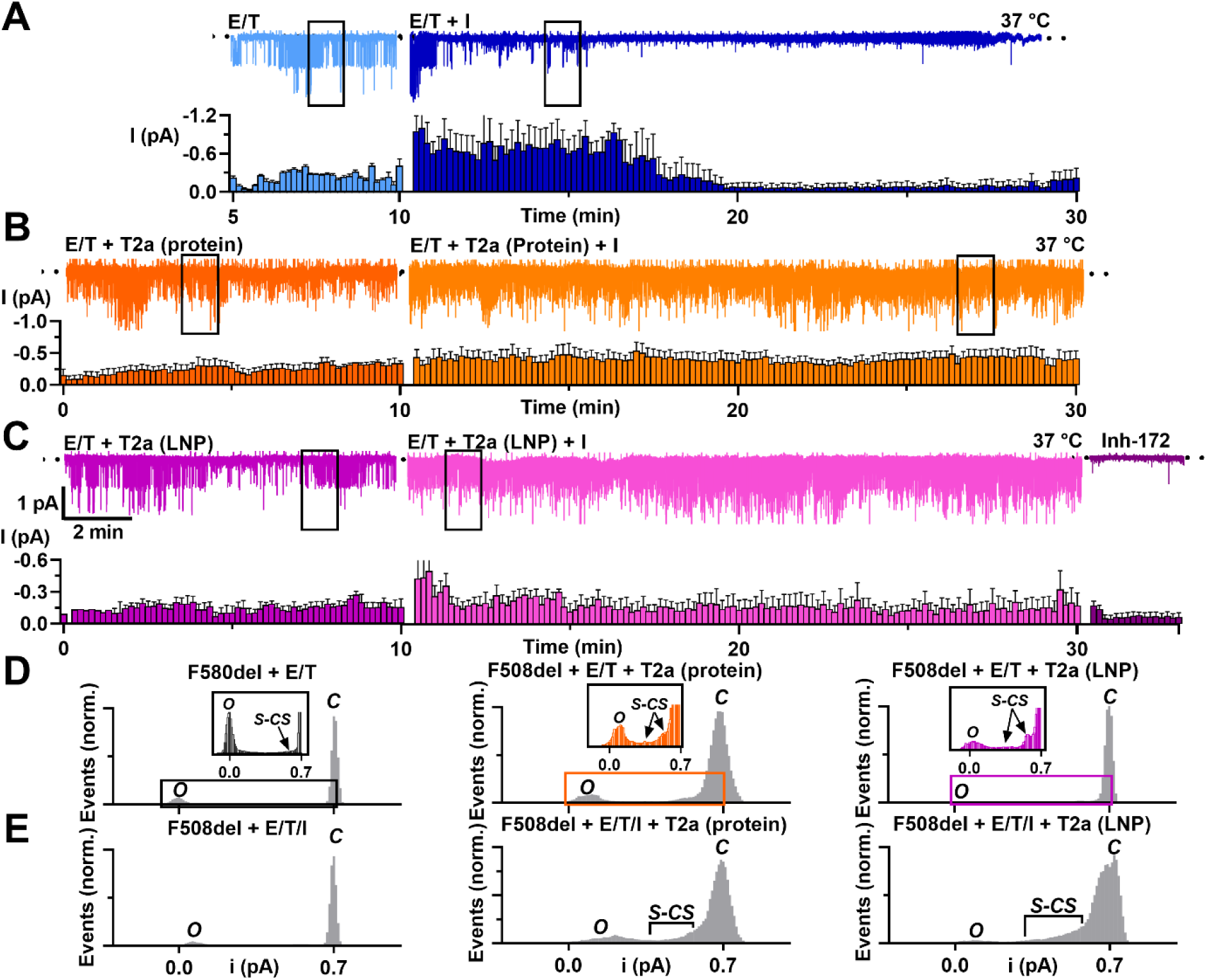
Nanobody T2a stabilizes F508del-CFTR channel gating in cell-free membrane patches. (A–C) Representative prolonged recordings (top) and summary total current (I) time courses (bottom) of F508del-CFTR Cl^−^ channels in excised inside-out membrane patches from BHK cells stably expressing F508del-CFTR. To increase the plasma membrane expression of F508del-CFTR, BHK cells were treated with elexacaftor (E, 2 μM) and tezacaftor (T, 3 μM) for 24 h at 37 °C prior to study. The recordings were made at 37 °C before and after ivacaftor (I, 1 μM) addition to the intracellular solution, which continuously contained ATP (1 mM) and PKA (75 nM). (A) Control recording. (B) Nanobody T2a protein (1 μM) was acutely added to the intracellular solution following channel activation at 25 °C prior to raising the temperature to 37 °C and commencing the recording. (C) Recording from a membrane patch excised from a F508del-CFTR-expressing BHK cell treated with T2a nanobody mRNA LNPs (2 ng μl^−1^) for 24 h at 37 °C after channel activation at 25 °C prior to raising the temperature to 37 °C and commencing the recording. Dotted lines indicate the closed channel state and downward deflections correspond to channel openings. Beneath the representative prolonged recordings, summary total current (I) time courses are shown. Data are means ± SEM (n = 4–5) with I values calculated for consecutive 10 s intervals; individual data points are omitted for illustration purposes. (D, E) Single-channel current amplitude histograms for the boxed 60-s portions of the recordings shown in A – C, after additional filtering at 50 Hz, either before (D) or after ivacaftor addition to the intracellular solution (E). In D, regions of the histograms are enlarged in the insets to show openings to S-CSs. Abbreviations, C, closed; O, open; S-CS, sub-conductance state.

**Figure S4.**
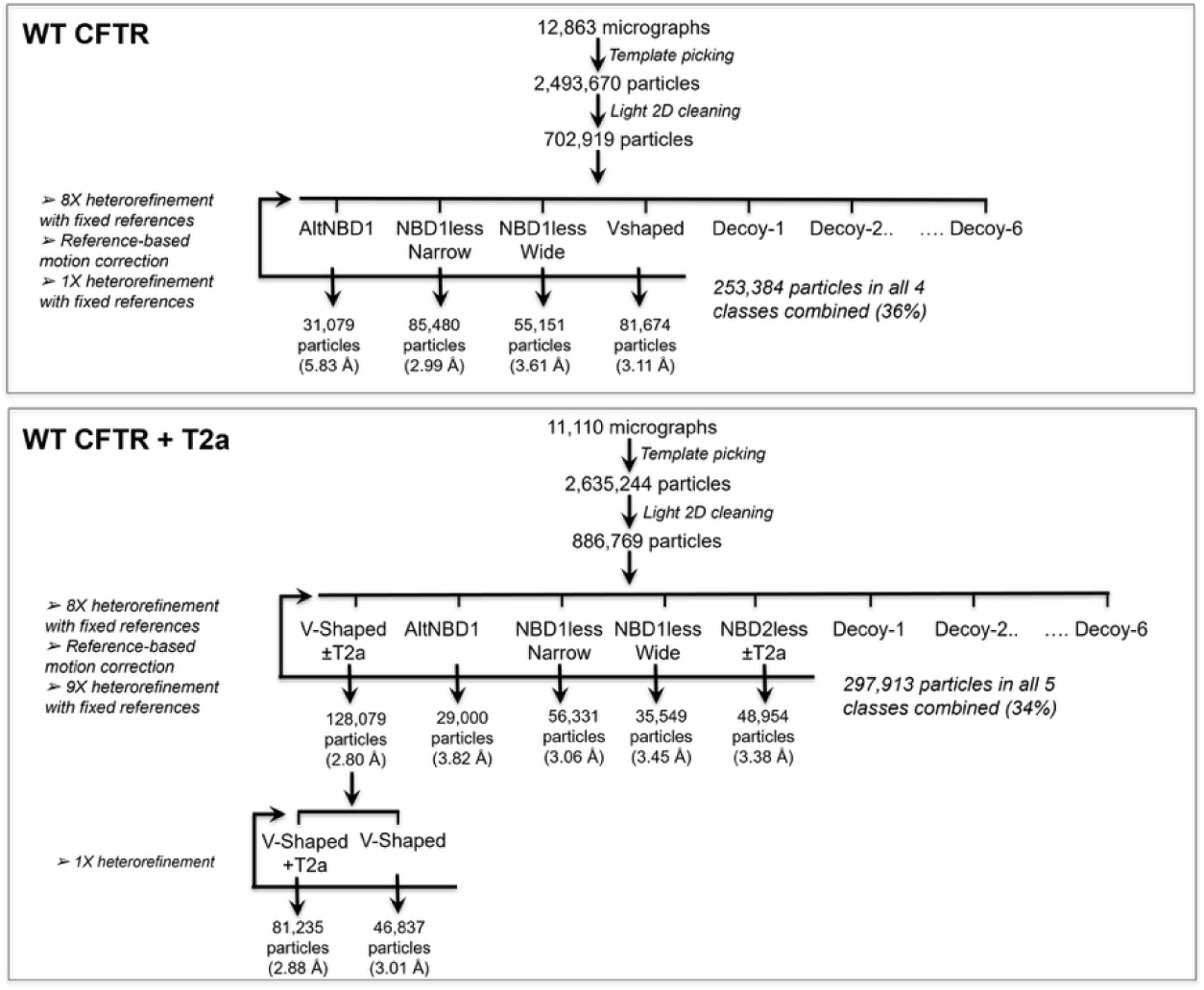
Cryo-EM classification flowchart. Schematic diagram illustrating the particle classification workflow used to process the cryo-EM datasets using cryoSPARC version 4.7.1. The V-Shaped+T2a and AltNBD1 structures presented in this paper come from the CFTR/T2a dataset, while the V-Shaped and NBD1lessNarrow structures come from the CFTR control dataset.

To analyze the single-channel behavior of F508del-CFTR treated with nanobody T2a, we used a similar approach to that adopted with WT-CFTR and observed openings to both the full open state and S-CSs (Figures 4D and E, S3B – D). By digitally filtering single-channel recordings at 50 Hz, we found that when F508del-CFTR was treated with nanobody T2a either acutely as purified protein or chronically with T2a LNPs, the frequency of openings to S-CSs was strongly increased. This effect was greatly enhanced by ivacaftor, with the result that discrete openings to S-CSs were no longer apparent in single-channel current amplitude histograms (Figures 4B – E, S3B – D). Although we observed openings to S-CSs of different amplitudes, we could not distinguish whether these represented openings of a single channel to different S-CSs ^37^ or S-CS openings of distinct channels because all membrane patches contained several active F508del-CFTR Cl^−^ channels. Nevertheless, the amplitude of S-CS openings was comparable between F508del-CFTR treated with T2a nanobody and untreated F508del-CFTR controls (Figures 4 and S3D). Similarly, the amplitude of the full open state of T2a nanobody-treated F508del-CFTR Cl^−^ channels was comparable to that observed in the absence of the nanobody (Figures 4B – D, S3D). Thus, nanobody binding leads to robust thermodynamic stabilization of F508del-CFTR, while permitting channel opening, albeit frequently to sub-conductance states. Given that the previously established interaction geometry of T2a with NBD1 blocks ATP-dependent NBD1-NBD2 association ^26^, these results suggest the existence of an active CFTR channel conformation without this association, which represents the canonical structural mechanism driving channel opening.

### Binding of T2a reveals a novel conformation of CFTR

We used cryo-EM to determine the structure of the complex between T2a and CFTR grown in the presence of lumacaftor. Purified human WT-CFTR was phosphorylated, concentrated to ∼8 µM, and incubated at 4 °C for 2 h with 2 mM ATP in the absence or presence of ∼24 µM purified T2a nanobody prior to flash-freezing and deposition on cryo-EM grids (see Methods). After data collection, micrographs were processed, and 3-dimensional reconstruction was performed as previously described using cryoSPARC ^38,39^ (Figure S4 and Table S1). In the absence of T2a, single-particle analysis produced a 3.1 Å reconstruction of CFTR in the “inverted V-shaped” topology where the NBDs are separated by about 20 Å (Figure 5A). This conformation is highly similar to the cryo-EM structure previously reported for dephosphorylated ATP-free CFTR ^40^, except for the lack of any density for residues in the R domain. The two models show less than 1.8 Å RMSD, with most of the differences observed at the level of NBD2, which was slightly rotated in our structure (Figure S5A). The TM helices align very well throughout the two structures. In contrast to published structures of the catalytically inactive E1371Q mutant ^25,41^ our cryo-EM analysis of WT-CFTR did not show a detectable population of NBD1-NBD2 associated structures likely because digitonin biases the energetic landscape away from this conformation.

**Figure 5.**
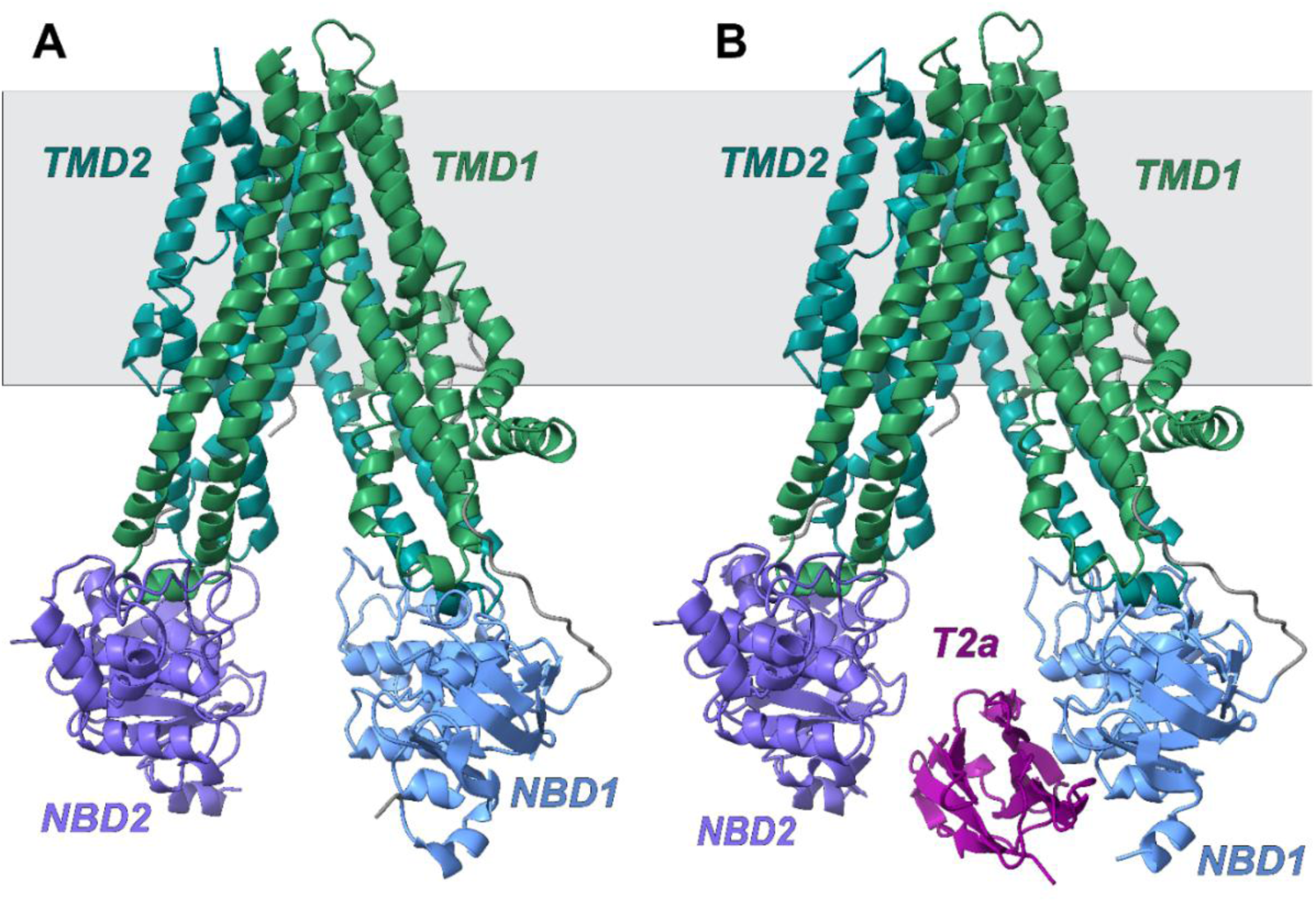
Cryo-EM structures of the “inverted V-shaped” conformation of CFTR in the absence and presence of nanobody T2a. (A) Structure of WT-CFTR from particles not bound to T2a. TMD1 (helices 1-6) is shown in green, TMD2 (helices 7-12) in teal with NBD1 in light blue and NBD2 in purple. (B) Structure of the major conformation of CFTR-T2a complex. The predicted location of the membrane is shown in grey.

The 2D classes from WT-CFTR samples incubated with the T2a nanobody clearly showed particles with density for both proteins. Classification methods identified five distinct CFTR conformations (Figure S4). The particles corresponding to the dominant inverted V-shaped conformation (comprising ∼1/3 of the population) were isolated using heterorefinement, and that population was further subdivided into particles with and without bound T2a using an additional round of heterorefinement. Non-uniform refinement of the inverted V-Shaped particles containing T2a, which comprised 27% of all particles in this WT-CFTR sample, yielded a map at 2.9 Å resolution, which enabled modelling of the complex (Figure S4). This T2a-bound inverted V-shaped structure (Figure 5B) shows the transmembrane helices in a conformation equivalent to that observed in the absence of T2a (RMSD < 1 Å) (Figure 5A). The structure of the nanobody was clearly visible, and the interface between T2a and NBD1 corresponded closely to that previously identified by X-ray crystallography (Figure S5B) ^26^. As a result, the nanobody was located between the two NBDs (Figure 5B), with a slight outward rearrangement of the NBDs when compared to the structure in the absence of nanobody (“apo” Figure S5C). This rigid-body displacement, which increased the distance between the NBDs by about 6 Å, was achieved by local rearrangements of the ICLs that connect NBD1 and NBD2 to the TM domains (Figure S5D).

**Figure S5.**
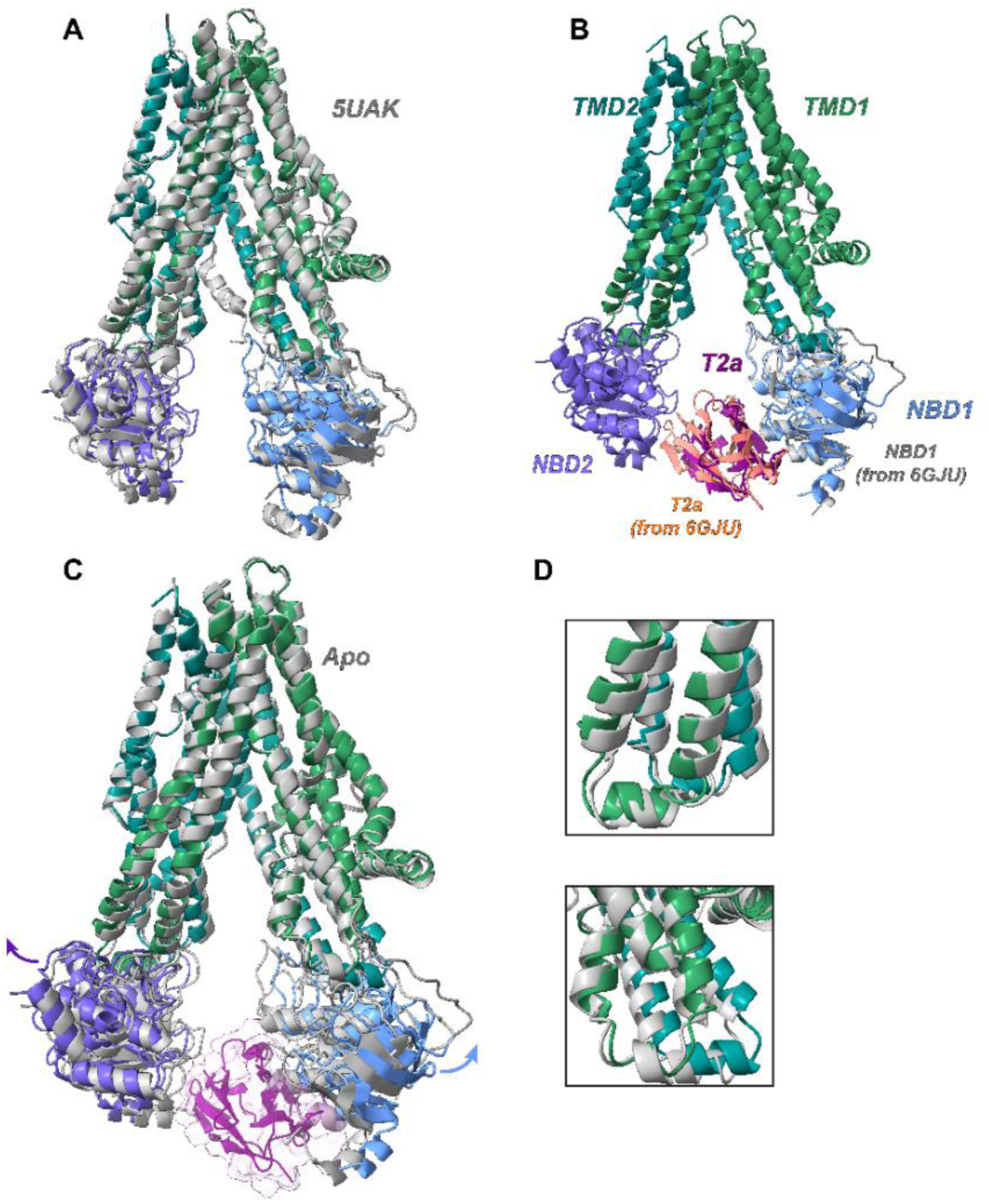
Structural analysis of the inverted V-shaped CFTR structures solved in the absence and presence of nanobody T2a. (A) Superimposition of the apo structure of CFTR (this work) onto the published structure of dephosphorylated ATP-free CFTR (PDB id: 5UAK, grey). (B) Superimposition of the NBD1-T2a complex previously obtained by crystallography (PDB id: 6GJU, grey and salmon) onto the inverted V-shaped conformation of the full-length CFTR bound to T2a (this work). (C) Superimposition of the inverted V-shaped structures in the absence (“apo”, grey) and presence (colored) of nanobody T2a, showing the slight motion of the NBDs (arrows). (D) Local rearrangement of the ICLs connecting the TMDs to NBD2 (top) and NBD1 (bottom), respectively.

Both in the absence and presence of T2a, the TM helices in the inverted V-shaped conformation meet towards the extracellular side of the membrane (Figure 5), preventing solvent access to the interior of the TM bundle and ion passage, which is in agreement with the analysis of the dephosphorylated ATP-free CFTR structure which concluded that this conformation corresponds to an inactive state ^40^.

We identified an additional conformation of the CFTR-T2a complex with a large rearrangement of NBD1 and the intracellular region of TMD1 (Figures 6 and S6). While the 2D classes showed molecular details characteristic of high-resolution images, the preferential orientation of these particles complicated 3D reconstruction. Nonetheless, iterative deconvolution of the anisotropic reconstructions using Ardecon (see Methods) yielded a 3.9 Å map in the consensus region. This map was directly interpretable in the TM region and NBD2, while reference atomic models for NBD1 and T2a nanobody could be unambiguously oriented in the lower-resolution regions of the map. This approach enabled an atomic model for this complete CFTR-T2a complex to be built and refined (Figure 6A). In this structure, NBD2 is bound to TMD2 in an equivalent manner to the canonical conformations of CFTR. In contrast, the NBD1-T2a complex is detached from the ICLs of TMD1 and docked instead to the C-terminus of the lasso motif ^40^ adjacent to the intracellular surface of the membrane (Figure S6A and B). The transmembrane region of this conformation is similar to that observed in the phosphorylated ATP-bound, NBD1:NBD2-dimerized structure previously reported for the hydrolysis-deficient E1371Q mutant of human CFTR ^41^, except for the absence of density for the intracellular regions of TM10 and TM11 and the coupling helix (ICL4) that connects them (Figure S6E). A portion of the volume usually occupied by these two segments showed density for a short α-helix likely to be derived from the C-terminus of the R domain (based on the sidechain density in the regions and observation of a continuous connection to the C-terminus of TMD2).

**Figure 6.**
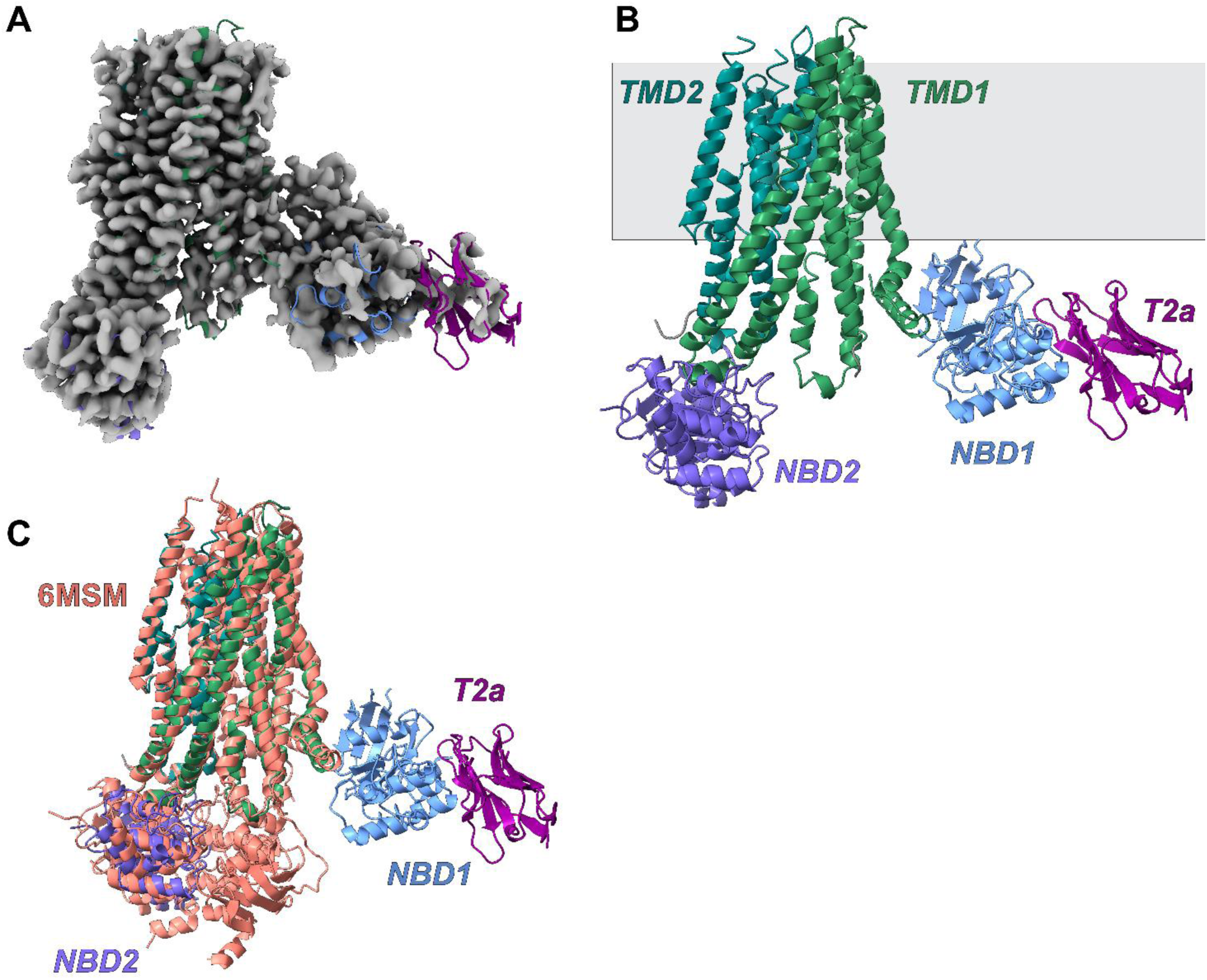
A novel structure of CFTR revealed by T2a nanobody binding. (A) EM density for the minor conformation of the CFTR-T2a complex. (B) Atomic model of the novel conformation. The predicted position of the membrane is shown in grey. (C) Structural comparison with the published phosphorylated ATP - bound structure of human CFTR (PDB id: 6MSM, salmon).

**Figure S6.**
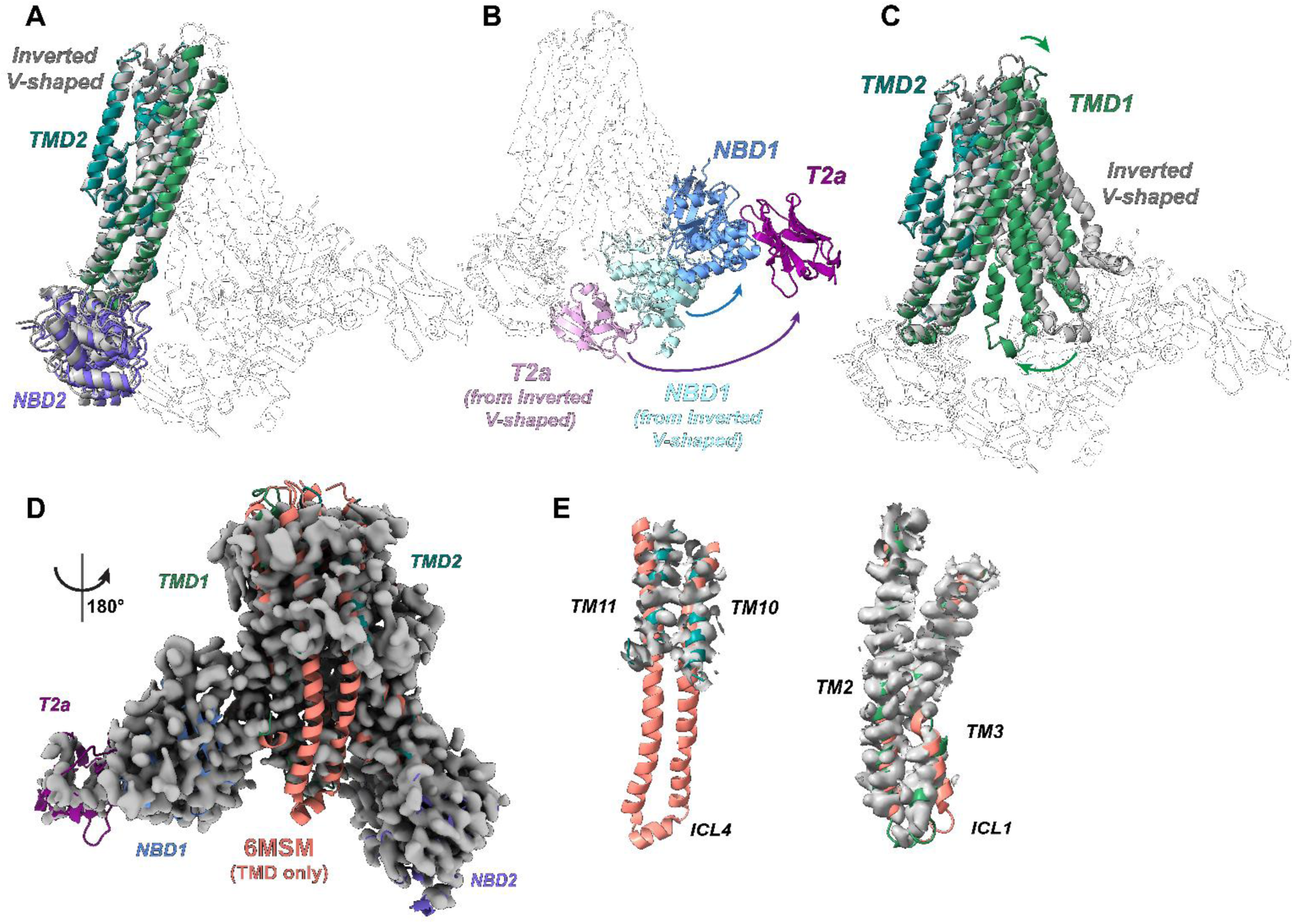
Structural similarities and conformational changes between the inverted V-shaped and the novel alternative conformation of the CFTR-T2a complex. (A) Structural alignment of NBD2-TMD2 (helices 4, 5, 7, 8, 9, 12) between the inverted V-shaped (gray) and novel (colored) conformations. For clarity the rest of the protein is represented in transparent ribbons. (B) Using the alignment in (A), a major conformational rearrangement of NBD1 (blue arrow) occurs which disengages it from the TMDs and reorients it towards the intracellular surface of the membrane. This motion requires a very large rotation, which is evident from the reorientation of the T2a nanobody (purple arrow) in comparison to the inverted V-shaped structure. For clarity, the rest of the protein is represented in transparent ribbons. (C) Superimposition of TMD helices from the two T2a-bound structures showing a reorganization of the two halves of the TM bundle, with the helices coming together on the intracellular side and moving slightly apart on the extracellular side. For clarity the rest of the protein is represented in transparent ribbons. (D) Superimposition of the published phosphorylated ATP-bound human CFTR structure (PDB id: 6MSM, salmon) on the novel CFTR-T2a state and its EM density. The NBDs of the published structure have been removed from its molecular model for clarity. (E) Zoomed-in view showing local comparisons of the TM2-ICL1-TM3 (right) and TM10-ICL4-TM11 (left) regions in the superposition in panel D (salmon from 6MSM, green from the alternative conformation of the CFTR-T2a complex).

The position of NBD1 in the novel conformation involves a large-scale translation and rotation compared to the inverted V-shaped structure of CFTR. NBD1 reorients towards the membrane, rotating along two orthogonal axes (Figure S6B). This movement disrupts all of the standard interactions between NBD1 and ICL1 or ICL4, which, in turn, show relatively weak density (Figure S6E). The N-terminal antiparallel β-sheet in the ATP-binding core subdomain of NBD1 instead binds to the C-terminal α-helix in the Lasso motif, which is uniquely found in ABCC subfamily of the ABC Superfamily proteins and comprises the first 68 residues in CFTR. While the EM density locally in this region is too poor to resolve a detailed view of this new interface, the overall binding geometry is clear. The C-terminal α-helix in the Lasso motif has previously been shown to be strictly required for CFTR expression ^42^ and function ^43^ and contains several rare CF-causing mutations.

The TM domain in the alternative CFTR-T2a complex adopts a very similar conformation to that of the phosphorylated ATP-bound structure of human CFTR (Figure 6C), where the helices in TMD1 and TMD2 move much closer to one another on the intracellular side of the membrane, while moving slightly further away from one another on the extracellular side (Figure S6C). These movements result in the formation of the internal chloride-conductance channel, which is not present in the inverted V-Shaped conformation of CFTR due to the details of the interhelical packing interactions in that conformation. Analysis of the structure of phosphorylated ATP-bound human CFTR revealed an extended Cl^−^ permeation pathway, and while a fully open channel was not observed, it was proposed that this CFTR conformation was close to that of an active state ^41^. When this structure is compared to the alternative CFTR-T2a complex, we observed that, except for the absence of density for the intracellular regions of TM10 and TM11, as mentioned above, they have very similar conformations, with an overall RMSD ∼2 Å. Modest shifts are observed in the positions of ICL1 and the C-terminal α-helix in the Lasso in the cytosolic region of the protein and TM9, TM10, and TM11 near the extracellular surface of the protein (Figure S6D and S6E). A larger shift is observed in this region at the N-terminus of TM12, and the first two turns at the N-terminus of TM8 become disordered. Although the limited resolution of the alternative CFTR-T2a complex structure prevents reliable modeling of the channel pore, the similar organization of TM helices in the phosphorylated ATP-bound inverted V-Shaped structure suggests the formation of a similar Cl^−^ permeation pathway. We therefore hypothesize that this novel CFTR-T2a complex represents an active conformation.

The novel conformation of the CFTR-T2a complex, which features an undocked NBD1, has not been reported previously. The observation of this complex raises the possibility of whether the alternative binding of NBD1 is induced by nanobody binding or can occur independently in native CFTR. To address this, we performed a complete cryo-EM analysis of a WT-CFTR sample prepared under identical conditions to those used for the CFTR-T2a complex, but in the absence of nanobody. Careful analysis revealed a diversity of conformations, including populations exhibiting an undocked NBD1. Most of these particles showed no visible density for NBD1, although, importantly, protein degradation was not observed on SDS-PAGE analysis (data not shown). However, a sub-population of the particles displayed density for NBD1 with a spatial arrangement closely resembling that observed in the alternative CFTR-T2a complex structure. Although the number of WT-CFTR particles was insufficient for high-resolution 3D reconstruction, 2D class averages clearly resolved NBD1 in a detached position relative to TMD1, matching the orientation seen in the CFTR-T2a complex (Figure S7A), while the TM helices adopted a similar organization to those observed in the nanobody-bound complex (Figure S7A). The WT-CFTR particles without visible NBD1 density were sufficiently abundant to build a 2.9 Å 3D map, enabling construction of a reliable atomic model (Figure S7B). Figure S7C shows that all other domains of CFTR are clearly present with the notable exceptions of the cytoplasmic ends of the TM10-ICL4-TM11 segments. Remarkably, there is a pore-like structure at the center of the TM helices that is indistinguishable from that observed in the CFTR-T2a complex (Figure S7D). This result demonstrates that NBD1 undocking is an intrinsic feature of CFTR dynamics, which facilitates the formation of a channel pore in the TMDs despite ICL4 being disordered.

**Figure S7.**
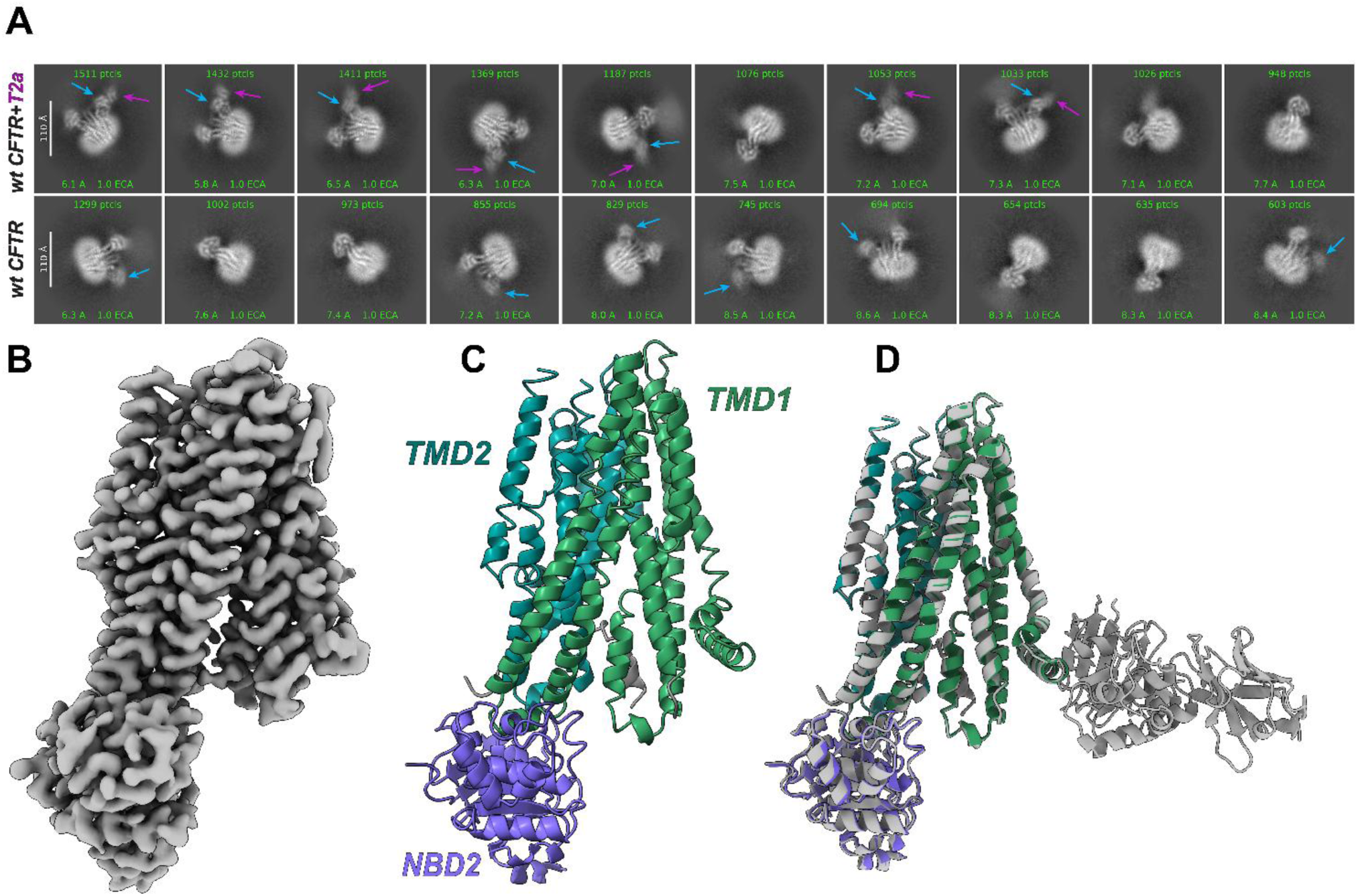
2D classes show that the undocked conformation of WT-CFTR is observed in the absence of T2a nanobody. (A) 2D classes showing the alternative NBD1 binding geometry in WT-CFTR cryo-EM datasets collected in the presence (top panel) or absence (bottom panel) of the T2a nanobody. The images show the 10 highest-population classes out of 50 produced by applying the standard 2D classification algorithm in cryoSPARC to the final particle stack for the undocked conformation generated by iterative multi-class heterorefinement applied to all of the major conformational classes in each dataset. The location of NBD1 and nanobody T2a are highlighted by blue and purple arrows, respectively. (B, C) EM density and atomic model of WT-CFTR lacking NBD1. (D) Superimposition of the CFTR-T2a complex with that of WT-CFTR lacking NBD1 showing a very similar arrangement of the TMDs.

## DISCUSSION

CFTR is expected to adopt a discrete number of conformations, where transition from a closed to open state of its chloride-conductance channel requires heterodimerization of the NBDs to drive rearrangement of the TMDs into a pore-like structure ^28,41^. The accepted gating model also assumes that, like other ABC proteins, the NBDs remain attached to the TMDs via the ICLs throughout the conformational cycle ^40^.

Our study indicates that other CFTR conformations may lead to channel opening, in which NBD1 detachment from TMD1 enables channel formation in the absence of NBD heterodimerization. We previously proposed that NBD1 detachment occurs naturally, based on labeling of CFTR-expressing cells by nanobodies whose epitope (which includes F508) is buried in the NBD1-TM1 interface ^26^ and thus requires NBD1 displacement for epitope unmasking, consistent with recent models supporting a dynamic NBD1-TM1 interface ^44^.

A consequence of NBD1 detachment is the rearrangement of the TM helices into a pore-like structure (Figure 6C) similar to that of phosphorylated ATP-bound CFTR ^41^ which is expected to be close to an ion-conducting state. Displacement of NBD1 is associated with disordering of the TM10-ICL4-TM11 segments and conformational rearrangements in TM2-ICL1-TM3 segment as well as in the extracellular regions of TM8 through TM12 at the opposite end of the protein (Figure S6D and E). Our structural data (Figs. 5-6 and S4-S8) and electrophysiological data (Figs. 3-4 and S3) together strongly support this novel CFTR conformation mediates the chloride-channel conductance observed from F508del-CFTR and WT-CFTR in cells treated with T2a LNPs. Conductance likely occurs through a dynamic pore sampling a combination of subconductance states (S-CSs) and fully open states (Fig. S3B-D), although the latter only form transiently after long-term exposure to T2a (Fig. S3A).

Deletion of F508 is likely to facilitate NBD1 detachment from TMD1 through a combination of domain destabilization and weakening of the NBD1-TMD1 interface, which would explain why F508del-CFTR opened to sub-conductance states when nanobody T2a was added acutely, whereas WT-CFTR closed for prolonged periods (Figures 3 and 4). However, WT-CFTR frequently opened to sub-conductance states when cells were treated with T2a mRNA LNPs. This result suggests that, when CFTR is exposed to the nanobody during its biosynthesis, binding of T2a might promote long-lasting NBD1 detachment.

Spontaneous NBD1 undocking in WT-CFTR is also supported by the presence of NBD1-less particles in our cryo-EM analyses of that protein, although the relative proportions observed in detergent micelles may not correspond to those existing in biological membranes. Interestingly, cryo-EM structures of ABC proteins with incomplete (or missing) NBDs have been reported previously, including the bile salt export pump (BSEP/ABCB11) ^45^ and multidrug resistance protein 4 (MRP4/ABCC4) ^46^. For MRP4, which belongs to the ABCC subfamily like CFTR, loss of NBD1 density was observed in all ligand-bound states ^46^. The apparent NBD1 detachment in these structures was also accompanied by a reorganization of the TM helices, reminiscent of that observed in CFTR (Figure S8).

Taken together, the data suggest that, in ABCC proteins (and potentially other ABC transporters), NBD1 detachment is a conserved feature enabling TMD reorganization that is linked to protein function, such as substrate recognition (e.g. MRP4^46^) or channel opening (CFTR), thus expanding current models of ABC protein regulation. The identification of CFTR conformations where NBD1 is detached from the TMDs but docked onto the C-terminal α-helix in the Lasso motif, a conserved structural feature in the ABCC family, further supports an evolved mechanism exploiting NBD1 detachment, and additional research will be needed to investigate how the possible interaction between NBD1 and the membrane may modulate this conformational equilibrium.

**Figure S8:**
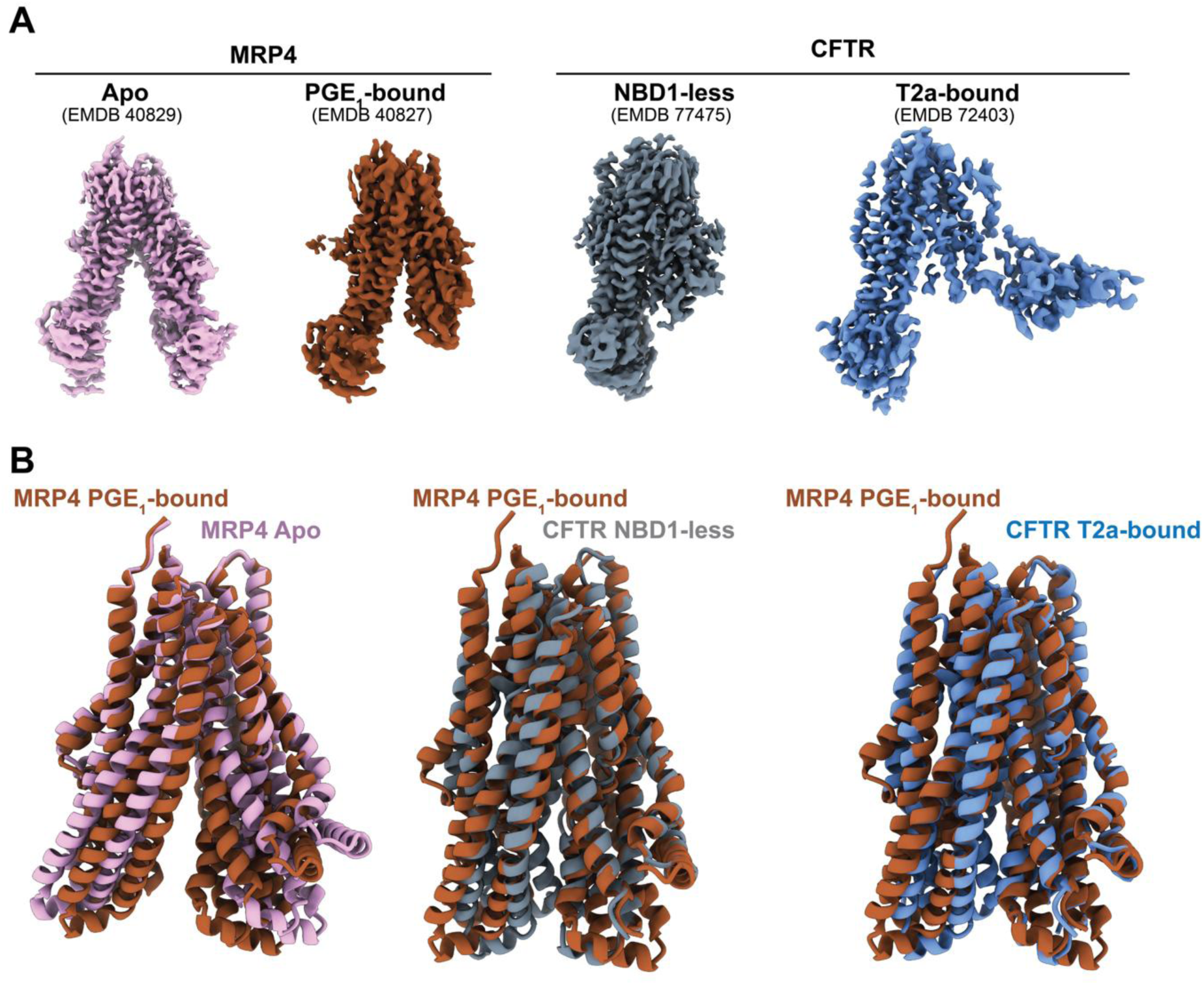
Absence of NBD1 density in ABCC proteins is associated with re-organization of the TM helices. A) Selected cryo-EM maps of human MRP4 ^46^ and CFTR (this study). For MRP4, the map shows all stably folded domains in the apo condition (pink) but lacks density for NBD1 when the PGE_1_ ligand is bound (brown). (B) Superimposition of the TMDs of MRP4 and CFTR. Left, MRP4 in the ligand-bound and apo states; middle, ligand-bound MRP4 and CFTR with undocked NBD1 (grey), and right, ligand-bound MRP4 and CFTR with bound T2a nanobody (blue).

Our data demonstrate that providing NBD1-stabilizing molecular chaperones in addition to the approved correctors strongly improves mutant CFTR function and supports the possibility that this approach can be used to augment current CF therapies. Therefore, using our target-specific nanobodies as tailored molecular chaperones opens a complementary therapeutic avenue for people with CF where pharmacological correction remains partial or the drugs are poorly tolerated ^16–18,21,47,48^.

As discussed above, the structural characterization enabled by our nanobodies expands the conformational landscape for CFTR, revealing that distinct conformational states can lead to channel opening, which could be critical to developing mutation-specific therapies. More broadly, we propose that developing tailored chaperones for other protein-trafficking diseases where pharmacological correction remains elusive offers a novel therapeutic strategy that can simultaneously advance our understanding of the molecular mechanisms underlying their pathophysiology.

**Table S1.**
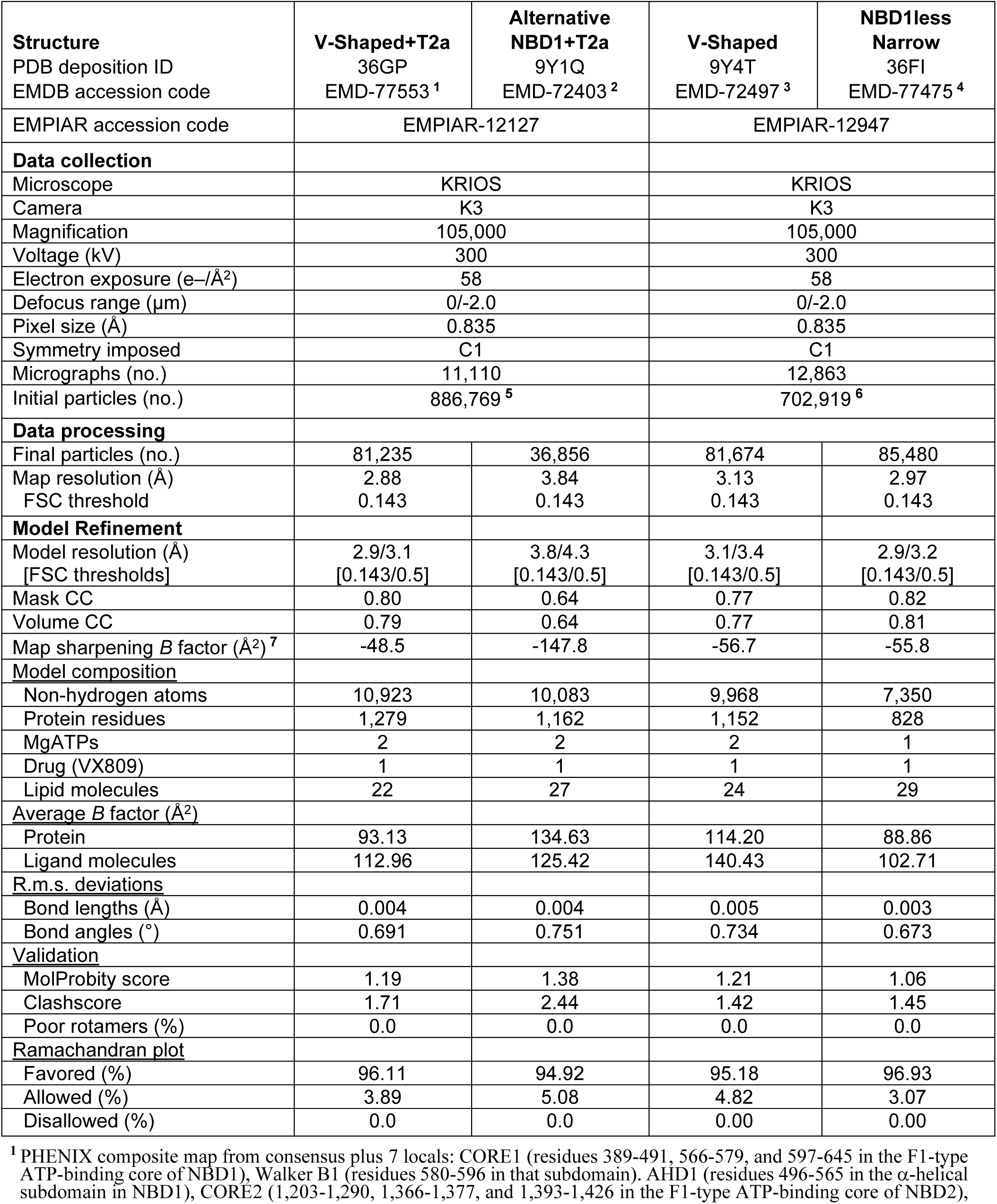
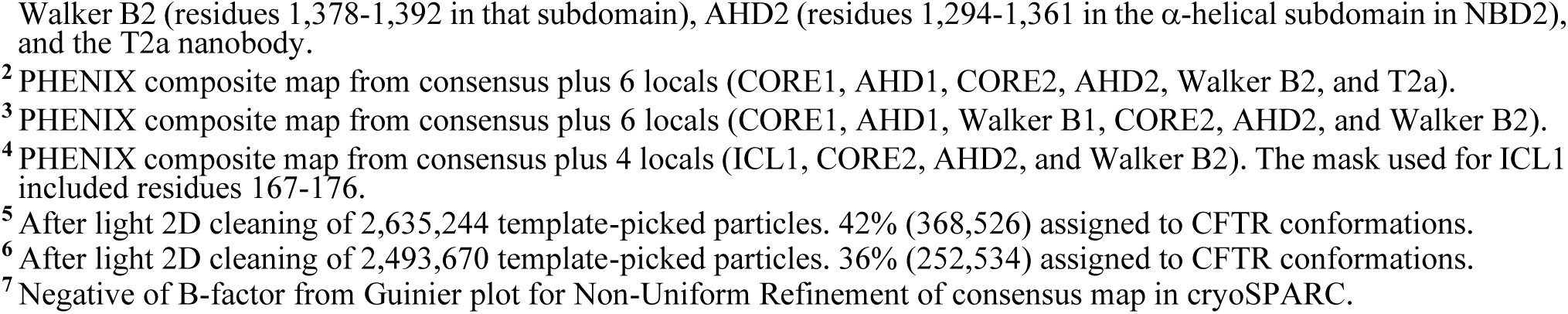
Cryo-EM data collection and refinement statistics for human WT-CFTR structures.

| Structure<br>PDB deposition ID<br>EMDB accession code | V-Shaped+T2a<br>36GP<br>EMD-77553<sup>1</sup> | Alternative<br>NBD1+T2a<br>9Y1Q<br>EMD-72403<sup>2</sup> | V-Shaped<br>9Y4T<br>EMD-72497<sup>3</sup> | NBD1less<br>Narrow<br>36FI<br>EMD-77475<sup>4</sup> |
| --- | --- | --- | --- | --- |
| EMPIAR accession code | EMPIAR-12127 |  | EMPIAR-12947 |  |
| **Data collection** |  |  |  |  |
| Microscope | KRIOS |  | KRIOS |  |
| Camera | K3 |  | K3 |  |
| Magnification | 105,000 |  | 105,000 |  |
| Voltage (kV) | 300 |  | 300 |  |
| Electron exposure (e-/Å<sup>2</sup>) | 58 |  | 58 |  |
| Defocus range (μm) | 0/-2.0 |  | 0/-2.0 |  |
| Pixel size (Å) | 0.835 |  | 0.835 |  |
| Symmetry imposed | C1 |  | C1 |  |
| Micrographs (no.) | 11,110 |  | 12,863 |  |
| Initial particles (no.) | 886,769<sup>5</sup> |  | 702,919<sup>6</sup> |  |
| **Data processing** |  |  |  |  |
| Final particles (no.) | 81,235 | 36,856 | 81,674 | 85,480 |
| Map resolution (Å) | 2.88 | 3.84 | 3.13 | 2.97 |
| FSC threshold | 0.143 | 0.143 | 0.143 | 0.143 |
| **Model Refinement** |  |  |  |  |
| Model resolution (Å)<br>[FSC thresholds] | 2.9/3.1<br>[0.143/0.5] | 3.8/4.3<br>[0.143/0.5] | 3.1/3.4<br>[0.143/0.5] | 2.9/3.2<br>[0.143/0.5] |
| Mask CC | 0.80 | 0.64 | 0.77 | 0.82 |
| Volume CC | 0.79 | 0.64 | 0.77 | 0.81 |
| Map sharpening *B* factor (Å<sup>2</sup>)<sup>7</sup> | -48.5 | -147.8 | -56.7 | -55.8 |
| **Model composition** |  |  |  |  |
| Non-hydrogen atoms | 10,923 | 10,083 | 9,968 | 7,350 |
| Protein residues | 1,279 | 1,162 | 1,152 | 828 |
| MgATPs | 2 | 2 | 2 | 1 |
| Drug (VX809) | 1 | 1 | 1 | 1 |
| Lipid molecules | 22 | 27 | 24 | 29 |
| **Average *B* factor (Å<sup>2</sup>)** |  |  |  |  |
| Protein | 93.13 | 134.63 | 114.20 | 88.86 |
| Ligand molecules | 112.96 | 125.42 | 140.43 | 102.71 |
| **R.m.s. deviations** |  |  |  |  |
| Bond lengths (Å) | 0.004 | 0.004 | 0.005 | 0.003 |
| Bond angles (°) | 0.691 | 0.751 | 0.734 | 0.673 |
| **Validation** |  |  |  |  |
| MolProbity score | 1.19 | 1.38 | 1.21 | 1.06 |
| Clashscore | 1.71 | 2.44 | 1.42 | 1.45 |
| Poor rotamers (%) | 0.0 | 0.0 | 0.0 | 0.0 |
| **Ramachandran plot** |  |  |  |  |
| Favored (%) | 96.11 | 94.92 | 95.18 | 96.93 |
| Allowed (%) | 3.89 | 5.08 | 4.82 | 3.07 |
| Disallowed (%) | 0.0 | 0.0 | 0.00 | 0.00 |

<sup>1</sup> PHENIX composite map from consensus plus 7 locals: CORE1 (residues 389-491, 566-579, and 597-645 in the F1-type ATP-binding core of NBD1), Walker B1 (residues 580-596 in that subdomain). AHD1 (residues 496-565 in the α-helical subdomain in NBD1), CORE2 (1,203-1,290, 1,366-1,377, and 1,393-1,426 in the F1-type ATP-binding core of NBD2),
Walker B2 (residues 1,378-1,392 in that subdomain), AHD2 (residues 1,294-1,361 in the α-helical subdomain in NBD2), and the T2a nanobody.
<sup>2</sup> PHENIX composite map from consensus plus 6 locals (CORE1, AHD1, CORE2, AHD2, Walker B2, and T2a).
<sup>3</sup> PHENIX composite map from consensus plus 6 locals (CORE1, AHD1, Walker B1, CORE2, AHD2, and Walker B2).
<sup>4</sup> PHENIX composite map from consensus plus 4 locals (ICL1, CORE2, AHD2, and Walker B2). The mask used for ICL1 included residues 167-176.
<sup>5</sup> After light 2D cleaning of 2,635,244 template-picked particles. 42% (368,526) assigned to CFTR conformations.
<sup>6</sup> After light 2D cleaning of 2,493,670 template-picked particles. 36% (252,534) assigned to CFTR conformations.
<sup>7</sup> Negative of B-factor from Guinier plot for Non-Uniform Refinement of consensus map in cryoSPARC.

## METHODS

### Cell culture

HEK293 cells stably expressing WT- or F508del-CFTR with 3 human influenza hemagglutinin (HA)-tags in ECL4 ^49^ were purchased from the Leuven Viral Vector Core. HEK293 cells cultured in DMEM, high glucose, GlutaMAX supplement (Gibco) supplemented with 6% FBS (v v^−1^) were maintained under puromycin selection at a concentration of 1 µg ml^−1^. For the HS-YFP assay, CFBE41o^−^ cells stably co-expressing F508del-CFTR and HS-YFP ^33^ were generously provided by Dr. N. Pedemonte (IRCCS Istituto Giannina Gaslini). CFBE41o^−^ expressing F508del-CFTR and HS-YFP cultured in MEM (Gibco) supplemented with 10% FBS (v v^−1^) and 2 mM L-glutamine were maintained under selection with 2 µg ml^−1^ puromycin and 0.75 mg ml^−1^ geneticin. For Ussing chamber studies, CFBE41o^−^ stably expressing F508del-CFTR were kindly supplied by Dr. E. J. Sorscher (Emory University) ^50^. CFBE41o^−^expressing F508del-CFTR were expanded in MEM (Gibco) supplemented with 10% FBS (v v^−1^), 5% L-glutamine (v v^−1^), 100 U ml^−1^ penicillin, 100 U ml^−1^ streptomycin and maintained under puromycin selection at a concentration of 4 µg ml^−1^. BHK cells stably expressing WT- and F508del-CFTR were a generous gift of Dr. M.D. Amaral (University of Lisboa) ^51^. They were cultured in a 1:1 mixture of Dulbecco’s modified Eagle’s Medium and Ham’s F-12 nutrient medium (Gibco) supplemented with 5% v v^−1^ fetal calf serum, 100 U ml^−1^ penicillin and 100 µg ml^−1^ streptomycin under methotrexate selection at a concentration of 200 μg ml^−1^. For structural studies, CFTR protein was expressed in Chinese hamster ovary (CHO) cells. Briefly, the nucleotide sequence of wild-type human CFTR, comprising a C-terminal StrepII-tag was inserted into a lentiviral vector immediately downstream of the tetracycline (Tet)-responsive element (TRE2). This positioned CFTR upstream of an internal ribosome entry site (IRES) followed by an open reading frame comprising puromycin N-acetyl-transferase, T2A and enhanced green fluorescent protein (EGFP), as described earlier ^52^. CHO cells constitutively expressing the reverse Tet transactivator (rtTA) were transduced with the packaged CFTR lentiviral vector, and 24 h after treatment with doxycycline CHO cells expressing EGFP were isolated by live-cell FACS sorting. The FACS sorted cells (designated D1557.s) exhibited a stable phenotype over several weeks in culture, including high cell density in large-scale suspension culture and strong CFTR expression within 24 h after doxycycline treatment (2 μg ml^−1^). All cells were cultured at 37 °C in a humidified atmosphere of 5% CO_2_.

### Lipid nanoparticles (LNPs) encapsulated T2a mRNA formulation and characterization

T2a-encoding mRNA with N1-methylpseudouridine (m1ψ) modification, Cap-0 and Cap-1 post-transcriptional modification and a 120 a-long polyadenylated tail was purchased from Tebu-Bio (Le Perray en Yvelines, France). The T2a nanobody was fused with a C-terminal myc tag (T2a-myc) or an enhanced green fluorescent protein tag (T2a-EGFP). mRNA LNPs were formulated as previously described ^53^. Briefly, a lipid mixture composed of D-Lin-MC3-DMA, sitosterol, DMG-PEG 2000, DSPC was prepared in 100% ethanol at 5 mM and a 50:38.5:1.5:10 molar ratio ^29,30^ and mRNA was diluted in 50 mM citrate buffer at pH 4.0. The lipid and mRNA solutions were mixed using the microfluidic device NanoAssemblr Ignite (Precision Nanosystems, Cytiva, Vancouver, Canada) at a 1:3 ratio followed by mini dialysis for ≥ 90 min against 10% sucrose/PBS using a Slide-A-Lyzer G3 Dialysis Cassette with 10,000 Da molecular-weight cutoff (Thermo Scientific, Waltham, MA, USA). Hydrodynamic size, polydispersity index and zeta potential were measured with dynamic light scattering using a Zetasizer Nano-ZS (Malvern Instruments, Malvern, UK). Messenger RNA encapsulation efficiency was assayed using the Quant-it RiboGreen RNA Assay Kit (Invitrogen, Waltham, MA, USA) or QuantiFluor RNA System (Promega, Madison, WI, USA).

### Sodium dodecyl sulfate-polyacrylamide gel electrophoresis (SDS-PAGE) and immunoblotting

HEK293 or CFBE41o^−^ cells grown in 3.5 cm dishes were transfected with T2a-myc or T2a-EGFP mRNA LNPs in DMEM or MEM (Gibco) medium, respectively. Cells were incubated with T2a mRNA LNPs (0.1 to 10 ng µl^−1^) at least overnight at 37 °C concurrently with 3 µM elexacaftor, 18 µM tezacaftor and 3 µM ivacaftor (ETI) or the corresponding vehicle DMSO. HEK293 cells were detached with 250 µl EDTA (0.5 mM), while CFBE41o^−^ cells were detached with 250 µl ice-cold PBS (scraped) and re-suspended with 3.5 mL medium. 2 mL of cell samples were centrifuged for 5 min at 2,000 x g, 4 °C.

Cell pellets were re-suspended in buffer containing 50 mM Tris-HCl pH 8.0, 150 mM NaCl, 0.1% SDS, 1% Triton X-100, 1 mM PMSF, protease inhibitor cocktail and incubated on ice for at least 1 h with occasional vortexing. Cell lysates were centrifuged for 15 min at 16,000 x g and the protein concentration of the supernatant determined with the Pierce BCA protein assay kit (Thermo Scientific). Proteins were stained with Cy5 N-hydroxysuccinimide ester (Amersham QuickStain Protein Labeling Kit - Cytiva) as loading control.

Cell extracts were separated by SDS-PAGE on 7.5% polyacrylamide gels and transferred to nitrocellulose membranes (Bio-Rad, Hercules, CA, USA) for immunodetection. Blots were blocked with 5% bovine serum albumin in Tris-buffered saline with added 0.05% Tween-20 for 1 h. CFTR was detected using monoclonal antibody 596 (1:10,000, CFTR Antibodies Distribution Program of the Cystic Fibrosis Foundation and the University of North Carolina at Chapel Hill ^32^) and T2a-EGFP was detected using anti-GFP antibody (0.4 µg ml^−1^, Roche, Darmstadt, Germany). Antibody binding was detected with anti-mouse HRP conjugated antibody (0.2 µg ml^−1^, Millipore, Darmstadt, Germany). The chemiluminescence signal was visualized using Luminata Forte Western HRP substrate (Millipore), detected with ImageQuant 800 Fluor and analysed by densitometry using ImageQuantTL software (both from Cytiva, Amersham, UK). After subtraction of the background signal, band C intensity was normalized to that of non-transfected cells treated with ETI and the loading control (total protein staining). Statistical significance was calculated by one-way ANOVA implemented in Prism 9 (GraphPad Software, San Diego, CA, USA, https://www.graphpad.com).

### Flow cytometry

HEK293 cells grown in 3.5 cm dishes were transfected with T2a-myc or T2a-EGFP mRNA LNPs in DMEM medium (Gibco). Cells were incubated with T2a mRNA LNPs (0.1 to 10 ng µl^−1^) at least overnight at 37 °C concurrent with 3 µM elexacaftor, 18 µM tezacaftor and 3 µM ivacaftor (ETI) or DMSO. HEK293 cells were detached with 250 µl EDTA (0.5 mM) and re-suspended with 3.5 mL medium. 1.5 mL of cell sample was centrifuged in flow cytometer adapted tubes for 5 min at 200 x g, 4 °C.

Cell pellets were co-incubated with anti-HA tag antibody (2 µg ml^−1^, BioLegend, San Diego, CA, USA) and 4’,6-diamidino-2-phenylindole (DAPI, 2.5 µM, Invitrogen) to monitor cell permeabilization. Antibody binding was detected using anti-mouse Alexa Fluor 700 conjugated antibody (1.3 µg mL^−1^, Invitrogen). The EGFP signal (Ex. 525/50 nm; Em. 500-550 nm), Alexa Fluor 700 signal (Ex. 725/20 nm; Em. 715-735 nm) and the DAPI signal (Ex. 450/50 nm; Em. 425-475 nm) were detected with the Gallios flow cytometer (Beckman Coulter, Brea, CA, USA). The Alexa Fluor 700 signal was recorded after gating on cells negative for DAPI (non-permeabilized cells). Data (medians) were analyzed with Kaluza software (Beckman Coulter). Statistical significance was calculated using one-way ANOVA implemented in Prism 9 (GraphPad Software).

### Halide-sensitive yellow fluorescent protein (HS-YFP) quenching assay

CFBE41o^−^ cells stably co-expressing F508del-CFTR and HS-YFP were reverse-transfected with T2a mRNA LNPs (2 ng µl^−1^) in 96 well plates at least overnight at 37 °C concurrent with elexacaftor (3 µM) and tezacaftor (18 µM) (ET) or DMSO treatment. The day after, cells were stimulated with 10 µM forskolin and potentiated with 3 µM ivacaftor in 200 µl medium for at least 20 min. Fluorescence before and after injection of 5 µl of buffer containing 3 M NaI, 60 mM KI, 38 mM KH_2_PO_4_, 230 mM Na_2_HPO_4_*2H_2_O, 110 mM D-glucose was measured (Ex. 485 nm; Em. 535 nm) with a plate reader SpectraMax iD3 (Molecular Devices, San Jose, CA, USA). After subtraction of the baseline (cells without corrector treatment), fluorescence was normalized to that before injection and non-linear regression was obtained with Prism 9 (GraphPad Software). Using the equation of the curve, we obtained the fluorescence 2 seconds after injection. Statistical significance was calculated using one-way ANOVA implemented in Prism 9 (GraphPad Software).

### Ussing chamber measurements

Preceding treatment, CFBE41o^−^ cells were trypsinized and passaged at a 1:2 ratio, and puromycin was withdrawn from the medium. The following day, CFBE41o^−^ cells were again trypsinized and reverse transfected with a solution containing MEM and either T2a mRNA LNPs (20 ng µl^−1^) or the vehicle (10% sucrose/PBS). Following transfection, one million cells were seeded onto Snapwell filters (Corning, NY, USA). Transfected cultures were maintained as submerged monolayers for 2-3 days. Once transepithelial resistance (R_t_) values of ≥ 500 Ω cm^2^ with the EVOM^2^ epithelial Volt-Ohm-meter (World Precision Instruments, Sarasota, FL, USA) were recorded, CFBE41o^−^ monolayers were treated with either DMSO or elexacaftor (3 µM) and tezacaftor (18 µM) by addition to media bathing the basolateral membrane and incubated for 24 h at 37 °C before mounting in Ussing chambers ^54^.

Transepithelial short-circuit (I_sc_) measurements were performed using EasyMount Ussing chambers (Physiologic Instruments, Venice, FL, USA) and a chloride concentration gradient with voltage clamped at 0 mV as previously described ^55^. Briefly, the basolateral compartment was perfused with a solution containing: 145 mM NaCl, 0.4 mM KH_2_PO_4_, 1.6 mM K_2_HPO_4_, 5 mM D-glucose, 1 mM MgCl_2_, and 1.3 mM calcium gluconate. The apical compartment was perfused with a 5 mM Cl^−^ solution containing 5 mM NaCl, 140 mM sodium gluconate, 0.4 mM KH_2_PO_4_, 1.6 mM K_2_HPO_4_, 5 mM D-glucose, 1 mM MgSO_4_ and 8 mM calcium gluconate. After a 10-20 min equilibration period in the presence of amiloride (100 µM) added to the solution bathing the apical membrane, CFBE41o^−^ monolayers were subjected to apical and basolateral treatment with forskolin (10 µM) and IBMX (100 µM), followed by apical application of ivacaftor (5 µM). Finally, CFTR_inh_-172 (20 µM) was added apically to assess total ion transport through CFTR after activation with forskolin and IBMX and potentiation with ivacaftor; following their addition, all compounds were continuously present in the solution bathing the apical membrane. Statistical analysis was performed by One-Way ANOVA to compare cAMP-induced I_sc_ and CFTR_inh_-172-sensitive I_sc_ between different treatment groups.

### Patch-clamp experiments

Prior to study, BHK cells expressing F508del-CFTR were treated with elexacaftor (2 μM) and tezacaftor (3 μM) for 24 h at 37 °C, whereas BHK cells expressing WT-CFTR were untreated. Immediately before commencing single-channel recordings, F508del-CFTR-expressing cells were washed in drug-free intracellular solution to remove CFTR correctors. However, the maximum period cells were left in drug-free intracellular solution before study did not exceed 30 min. To test the action of the T2a nanobody on F508del-CFTR, the purified T2a nanobody (1 μM) was added to the intracellular solution bathing excised inside-out membrane patches from BHK cells expressing F508del-CFTR treated with elexacaftor and tezacaftor. Alternatively, F508del-CFTR-expressing BHK cells were transfected with T2a mRNA LNPs (2 ng μl^−1^) and then incubated with elexacaftor (2 μM) and tezacaftor (3 μM) for 24 h at 37 °C.

CFTR Cl^−^ channels were recorded in excised inside-out membrane patches from BHK cells heterologously expressing CFTR using an Axopatch 200B patch-clamp amplifier and pCLAMP software (version 10.4) both from Molecular Devices as described previously ^56^. The pipette (extracellular) solution contained 140 mM N-methyl-D-glucamine, 140 mM aspartic acid, 5 mM CaCl_2_, 2 mM MgSO_4_ and 10 mM N-tris[hydroxymethyl]methyl-2-aminoethanesulfonic acid (TES), adjusted to pH 7.3 with Tris ([Cl^−^], 10 mM). The bath (intracellular) solution contained 140 mM NMDG, 3 mM MgCl_2_, 1 mM CsEGTA and 10 mM TES, adjusted to pH 7.3 with HCl ([Cl^−^], 147 mM; free [Ca^2+^], < 10^−8^ M) and was maintained at 37 °C.

CFTR Cl^−^ channels were activated promptly within 5 min following membrane patch excision using the catalytic subunit of protein kinase A (PKA; 75 nM) and ATP (1 mM) before voltage was clamped at –50 mV. Wild-type CFTR was activated at 37 °C, whereas F508del-CFTR was activated at room temperature before temperature was raised to 37 °C once channel activation was complete. To test the effects of purified T2a nanobody protein on the CFTR Cl^−^ channel, we first recorded single-channel activity for 5 - 10 min in the presence of ATP (wild-type, 0.3 mM; F508del-CFTR, 1 mM) and PKA (75 nM) before adding T2a nanobody (1 μM) to the intracellular solution and acquiring further single-channel activity. For wild-type CFTR, we acquired prolonged recordings in the presence of nanobody T2a, whereas for F508del-CFTR, we acquired 5 - 10 min of single-channel data with nanobody T2a before adding ivacaftor (1 μM) to the intracellular solution and acquiring 30 - 40 min of single-channel data. For F508del-CFTR-expressing cells treated with T2a nanobody mRNA LNPs, following membrane patch excision and channel activation, temperature was raised to 37 °C and 5 - 10 min of single-channel data acquired before ivacaftor (1 μM) was added to the intracellular solution and single-channel data acquired for 30-40 min. To minimize channel rundown, PKA (75 nM) and ATP (0.3 or 1.0 mM) were added to all intracellular solutions. On completion of experiments, the recording chamber was thoroughly cleaned before reuse ^57^. In this study, we used excised inside-out membrane patches containing ≤ 5 active channels. To determine channel number, we used the maximum number of simultaneous channel openings observed during an experiment determined using conditions that robustly potentiate CFTR activity and recordings that were long enough to ascertain the correct number of channels ^58^.

Single-channel currents were acquired directly to computer hard disc after filtering at a corner frequency of 500 Hz using an eight-pole Bessel filter (model F-900C/9L8L, Frequency Devices Inc., Ottawa, IL) and digitizing at a sampling rate of 5 kHz using a Digidata 1440A (Molecular Devices) and pCLAMP software. To visualize openings to S-CSs, we digitally filtered single-channel data at 50 Hz using pCLAMP software. To measure single-channel current amplitude (i), either Gaussian distributions were fit to current amplitude histograms or cursors were used. For open probability (P_o_) measurements of WT-CFTR using data digitized at 5 kHz only, lists of open- and closed-times were generated using a half-amplitude crossing criterion for event detection and dwell time histograms constructed as previously described ^59^; transitions < 1 ms were excluded from the analysis [eight-pole Bessel filter rise time (T_10-90_) ∼0.73 ms at f_c_ = 500 Hz]. Histograms were fitted with one or more component exponential functions using the maximum likelihood method. For illustration purposes, single-channel records were filtered at 500 Hz, digitized at 5 kHz and, where indicated, digitally filtered at 50 Hz, before file size compression by either 5-fold or 500-fold data reduction using pCLAMP software.

Results are expressed as means ± SEM of n observations, where n represents the number of individual membrane patches obtained from different cells. Using SigmaPlot (version 13.0, Systat Software Inc., San Jose, CA), we tested for differences between 2 groups of data acquired within the same experiment with Student’s t-test. Differences were considered statistically significant when P < 0.05.

### Cryo-EM sample preparation

Methods were equivalent to those previously described for the structural characterization of G551D-6SS-CFTR ^25^ except for the addition of 1.5 µM Lumacaftor during cell culture at the same time that protein expression was induced by addition of 1 µg/ml of doxycycline. In brief, stable lentiviral transformants of CHO cells verified to produce high level expression of WT-CFTR ^60^ were solubilized in 0.5% dodecyl-β-D-maltoside (DDM), 0.1% cholesteryl hemisuccinate (CHS), 10% (v v^−1^) glycerol, 150 mM NaCl, 2 mM DTT, 0.2 mM TCEP, 20 mM HEPES pH 7.5, and Roche cOmplete EDTA-free Protease Inhibitors (Millipore-Sigma, St. Louis, MO, USA). The detergent extract was loaded onto a Strep-Tactin affinity column (IBA LifeSciences, Göttingen, DE), which bound CFTR via a C-terminal Twin-Strep-tag, and washed extensively with the same buffer containing 0.06% (w v^−1^) digitonin instead of DDM/CHS prior to elution with 4 mM *d*-desthiobiotin. Purification was completed by two sequential steps of gel-filtration on Superose 6 Increase columns in TNM buffer (0.06% (w v^−1^) digitonin, 200 mM NaCl, 3 mM MgCl_2_, 1 mM DTT, 50 mM Tris-Cl pH 7.5). The first was performed on a 10 mm ID column with 2 mM ATP and 10% (v v^−1^) glycerol in the buffer, while the second was performed on a 3.2 mm ID microbore column with 150 mM ATP and no glycerol in the buffer. The protein was tumbled at 4 °C with TEV protease for 4 h and then PKA for an additional hour prior to the first gel-filtration step. The CFTR monomer peak from the second gel filtration step was concentrated 2-5 fold to a final concentration of ∼1.5 mg ml^−1^ and then incubated on ice for 2 h with the T2a nanobody at an ∼3:1 ratio (T2a:CFTR) prior to deposition of a 3 µl aliquot on a cryo-EM grid using a Vitrobot Mark IV System (ThermoFisher, Waltham, MA). The T2a nanobody was purified as previously described ^26^ before transfer by gel filtration to TNM buffer without digitonin and concentrated to ∼3 mg ml^−1^ in an Amicon Ultra Centrifugal Filter 10 kDa MWCO (Millipore-Sigma) in a swinging-bucket rotor (14,000 x g). UltraAuFoil 300 mesh R 0.6/1 grids (Quantifoil, Großlöbichau, Germany) were treated for 25s in a Solarus Plasma Cleaner 950 (Gatan, Pleasanton, CA, USA) with O_2_/H_2_ flow-rates of 27.5/6.4 sccm and 15 W cleaning power. Sample deposition and blotting were performed at 4 °C and 100% humidity applying a blot-force of 2-4 with Whatman 1 filter paper (Whatman Inc., Piscataway, NJ, USA) for 6-10 s prior to plunging into liquid ethane.

### Cryo-EM data collection

Leginon was used to collect data on a 300 kV Titan Krios electron microscope (Thermo Scientific) with a K3 camera and imaging filter (Gatan) at the Columbia University Cryo-EM Center using counting mode. The 2.5 s exposures were dose-fractionated into 50 frames. Details are given in Table S1.

### Cryo-EM structure determination

Particle classification and 3-dimensional volume reconstructions were performed in cryoSPARC (Structura Biotechnology, Toronto, Canada) using our previously reported Global Conformational Ensemble Reconstruction methods ^61^. In brief, following an exhaustive search for alternative conformations including 3D variability analyses using the full molecular envelope, iterative heterorefinement against the final set of major CFTR conformations and six decoy volumes was performed as schematized in Figure S4 using an initial resolution of 6 Å, followed by *ab initio* reconstruction and then non-uniform refinement of the particles assigned to each conformational class. Following generation of consensus maps for each structure in cryoSPARC, local refinements were performed on the two NBDs separately or on NBD1 and T2a together for the structures containing T2a. The outputs from the initial local refinements were used as inputs for an additional round of local refinements in smaller masks covering internally rigid structures that rotate relative to one another (i.e., T2a, the ATP-binding core subdomain and the α-helical subdomain in both NBDs, the α-helix following the Walker B motif in NBD2, and, exclusively for the inverted V-Shaped structure, the equivalent α-helix in NBD1. Gaussian rotation and translation restraints in ranges from 0-4 degrees and 0-4 pixels, respectively, were used for all local refinements, with the magnitudes being determined empirically by trial-and-error based on inspection of the output density maps. Composite map and half-map generation were performed in PHENIX ^62^ using local weighting to a coordinate model after manual alignment of all local maps and half maps to the consensus full map in ChimeraX (UCSF, San Francisco, CA, USA)^63^. The coordinate model used for composite map generation was produced by one round of real-space refinement in PHENIX against a crude manual composite map made from the aligned consensus and local maps using the Volume Maximum function in ChimeraX. For the AltNBD1 map, the initial composite map from cryoSPARC was used as input for anisotropy deconvolution in ARDECON with parameters optimized as recommended ^64^, and the PHENIX model-sharpened map from that procedure was used to seed non-uniform refinement in cryoSPARC to generate a new consensus map. That updated consensus map, which showed somewhat reduced orientational bias compared to the original consensus map, was used for another round of local refinements in cryoSPARC and composite map generation in PHENIX. These sequential ARDECON, local refinement, and composite map generation procedures were repeated two additional times to produce the final AltNBD1 composite map (Figure 6 and Table S1). Manual model rebuilding was performed using COOT ^65^ based on previously published structures ^40,41^ and real-space refinement of coordinate models and model-validation calculations were performed in PHENIX.

### Data Presentation

Structural figures were generated using UCSF ChimeraX. Plots were generated using GraphPad Prism 9. Flow cytometry histogram overlays were generated with FlowJo (Waters Biosciences Software, Ashland, OR, USA https://flowjo.com/flowjo/download). All the figures were assembled using Adobe Illustrator (San Jose, CA, USA https://www.adobe.com/be_fr/products/illustrator).

## QUANTIFICATION AND STATISTICAL ANALYSIS

The quantification and statistical analyses are integral parts of the algorithms used. Details are described in the main text and methods sections.

## SUPPLEMENTAL DOCUMENT

### T2a mRNA sequence

ATGCAGGTGCAGCTGCAGGAGTCAGGAGGAGGACTGGTGCAGGCAGGAGGATCTCTGAGACTGTCTTGCGCCGCTAGCGGCAGCATCTTTAGGATCGACGCTATGGGCTGGTACAGGCAGGCCCCAGGAAAACAGAGAGAGCTGGTGGCTCACAGCACAAGCGGAGGCAGCACCGATTACGCCGATAGCGTGAAGGGCAGGTTCACCATCAGCCGGGACAACGCCAAGAACACCGTGTACCTGCAGATGAACAGCCTGAAGCCCGAGGACACCGCCGTGTACTATTGCAACGCCGACGTGCGAACCCGCTGGTACGCCAGCAACAACTACTGGGGACAGGGAACACAGGTCACCGTGTCTAGCGGAAGCGGATCT

## RESOURCE AVAILABILITY

### Lead contact

Requests regarding reagents and further information may be addressed to the lead contact, Cedric Govaerts.

### Materials availability

This study did not generate new unique reagents.

### Data and code availability

- The cryo-EM movies used to generate the structures reported in this paper have been deposited in EMPIAR (EMPIAR-12127 and EMPIAR-12947). All consensus, local, and composite maps have been deposited in the EMDB (EMD-71697, EMD-71698, EMD-77553, EMD-71709, EMD-71710, EMD-71711, EMD-71713, EMD-71714, EMD-71756, EMD-71757, EMD-71758, EMD-71759, EMD-71760, EMD-71761, EMD-71762, EMD-71763, EMD-71764, EMD-71765, EMD-72403, EMD-72491, EMD-72492, EMD-72493, EMD-72494, EMD-72495, EMD-72496, EMD-72356, EMD-72497, EMD-77475, EMD-77407, EMD-77409, EMD-77470, EMD-77471, EMD-77472, EMD-77473, EMD-77474, EMD-77492), and the corresponding coordinate models have been deposited in the PDB (36GP, 9Y1Q, 9Y4T, and 36FI) for public release at the time of publication.
- Any additional information required to analyze the data reported in this paper is available from the lead contact upon request.
- This paper does not report original code.

## AUTHOR CONTRIBUTIONS

M.O. performed the CFTR maturation experiments and the HS-YFP quenching assays and analyzed the data with assistance of C.G. M.O. provided nanobodies and LNPs samples for Ussing chamber, patch-clamp and cryo-EM experiments. T.R. performed Ussing chamber measurements and analyzed the data with assistance of A.B. J.N.C. and M.R. carried out patch-clamp experiments and analyzed the data with assistance of D.N.S. A.S.P., B.J.L. and Z.R performed cryo-EM experiments and analyzed the data with assistance of J.F.H. and C.G. Z.Y. prepared CFTR samples for cryo-EM experiments. J.C.K. provided stable cell line for CFTR expression. M.O., T.R., J.N.C., M.R., A.B., M.A.M., D.N.S., J.F.H., C.G. contributed to writing the manuscript. C.G. oversaw the project.

## ACKNOWLEDGMENTS

We thank A. des Rieux and F. Debuisson for their excellent assistance with LNPs formulation, J. Baranwal and L. Prasad for assistance generating composite cryoEM maps and refining the coordinate models, and Kathrin Seidel and Marika Drescher for expert technical support with cell culture. This work was supported by (i) C.G.: the Fonds Forton, the Welbio (grant CR-2022), the Association luxembourgeoise de la lutte contre la mucoviscidose, the CF Trust, the Fondation Air Liquide and the Fondation ULB; (ii) D.N.S: the CF Trust (SRC 021); (iii) J.F.H. is grateful for 25 years of financial support from the US Cystic Fibrosis Foundation; (iv) J.C.K.: the Cystic Fibrosis Foundation (KAPPES18XX0, KAPPES20XX0) and (v) M.A.M.: the German Research Foundation (CRC 1449 – 431232613, 450557679 to M.A.M.) and the German Federal Ministry of Education and Research (82DZL009C1 and 01GL2401A to M.A.M.). C.G. is a senior Research Associate of the FRS-FNRS. J.N.C. is a CF Trust-supported PhD student (SRC 024).

